# Class A GPCRs use the membrane potential to increase their sensitivity and selectivity

**DOI:** 10.1101/587576

**Authors:** Daria N. Shalaeva, Dmitry A. Cherepanov, Michael Y. Galperin, Gert Vriend, Armen Y. Mulkidjanian

## Abstract

The human genome contains about 700 genes of G protein-coupled receptors (GPCRs) of class A; these seven-helical membrane proteins are the targets of almost half of all known drugs. In the middle of the helix bundle, crystal structures revealed a highly conserved sodium-binding site, which is connected with the extracellular side by a water-filled tunnel. Sodium ions are observed in GPCRs crystallized in their inactive conformations, but not in GPCRs that were trapped in agonist-bound active conformations. The escape route of the sodium ion upon the inactive-to-active transition and its very direction, either into the cytoplasm or back outside the cell, hitherto remained obscure. We modeled sodium-binding GPCRs as electrogenic carriers of sodium ions. In this model the sodium gradient over the cell membrane would increase the sensitivity of GPCRs if their activation is thermodynamically coupled to the translocation of the sodium ion into the cytoplasm, while decreasing it if the sodium ion retreats into the extracellular space upon receptor activation. The model quantitatively describes the available data on both activation and suppression of distinct GPCRs by membrane voltage. The model also predicts selective amplification of the signal from (endogenous) agonists if only they, but not their (partial) analogs, could induce sodium translocation. Comparative structure and sequence analyses of sodium-binding GPCRs indicate a key role for the conserved leucine residue in the second transmembrane helix (Leu2.46) in coupling sodium translocation to receptor activation. Hence, class A GPCRs appear to utilize the energy of the transmembrane sodium potential to increase their sensitivity and selectivity.

## Introduction

G-protein coupled receptors (GPCRs) are integral membrane proteins that consist of seven transmembrane (TM) helices surrounding a relatively polar core [1–4] (Fig. 1A-C). In most GPCRs, binding of the endogenous ligand (agonist) by the "inactive" form of protein causes a conformational change of the helical bundle (Fig. 1). The activated GPCR interacts with a G protein, which then triggers the intracellular signaling cascade. GPCRs are divided into several classes: rhodopsin-like receptors (class A), secretin receptors (class B), glutamate receptors (class C), fungal mating pheromone receptors (class D), cAMP receptors (class E), and frizzled receptors (class F). Sequence and structure comparisons of GPCRs support the notion that most, if not all, of them have a common evolutionary origin [5–7].

GPCRs are widespread among eukaryotes and are intensively studied for their ability to regulate various cellular processes. Different classes of GPCRs are unevenly represented in sequenced genomes from various eukaryotic lineages. In humans, class A GPCRs are the largest protein family with about 700 genes, whereas all other classes of GPCRs together have only about 150 representatives [6]. Human GPCRs serve as targets for about half of all known drugs [1].

The recent avalanche of high-resolution X-ray structures of GPCRs in ligand-free, agonist-bound, and antagonist-bound forms revealed many important features of their functioning [4, 8–16]. The major feature is that activation of GPCRs is associated with a large displacement of the cytoplasmic half of helix 6. A conserved Trp residue in the middle of this helix (Trp6.48 according to the Ballesteros–Weinstein numeration for class A GPCRs as modified by Isberg et al. [17, 18], used hereafter) serves as the pivot of this motion, see Fig. 1C and [2, 9, 19–25]. Another feature uncovered by these structures was the presence of Na^+^ ions in the vicinity of the ligand-binding sites in several class A GPCRs [12, 15, 16, 26–28]. The Na^+^ binding site is connected to the extracellular side through a clearly visible tunnel, but separated from the cytoplasmic side by a cluster of hydrophobic residues (Fig. 1A, B and [25, 26]). Analysis of the crystal structures of GPCRs in the active state (bound with agonists) shows that the Na^+^-binding pocket collapses from ~200 to 70 Å^3^ due to the movements of TM helices upon activation [26]. There is no space for the Na^+^ ion in the active state, suggesting that it leaves the receptor upon activation [12].

**Fig. 1.**
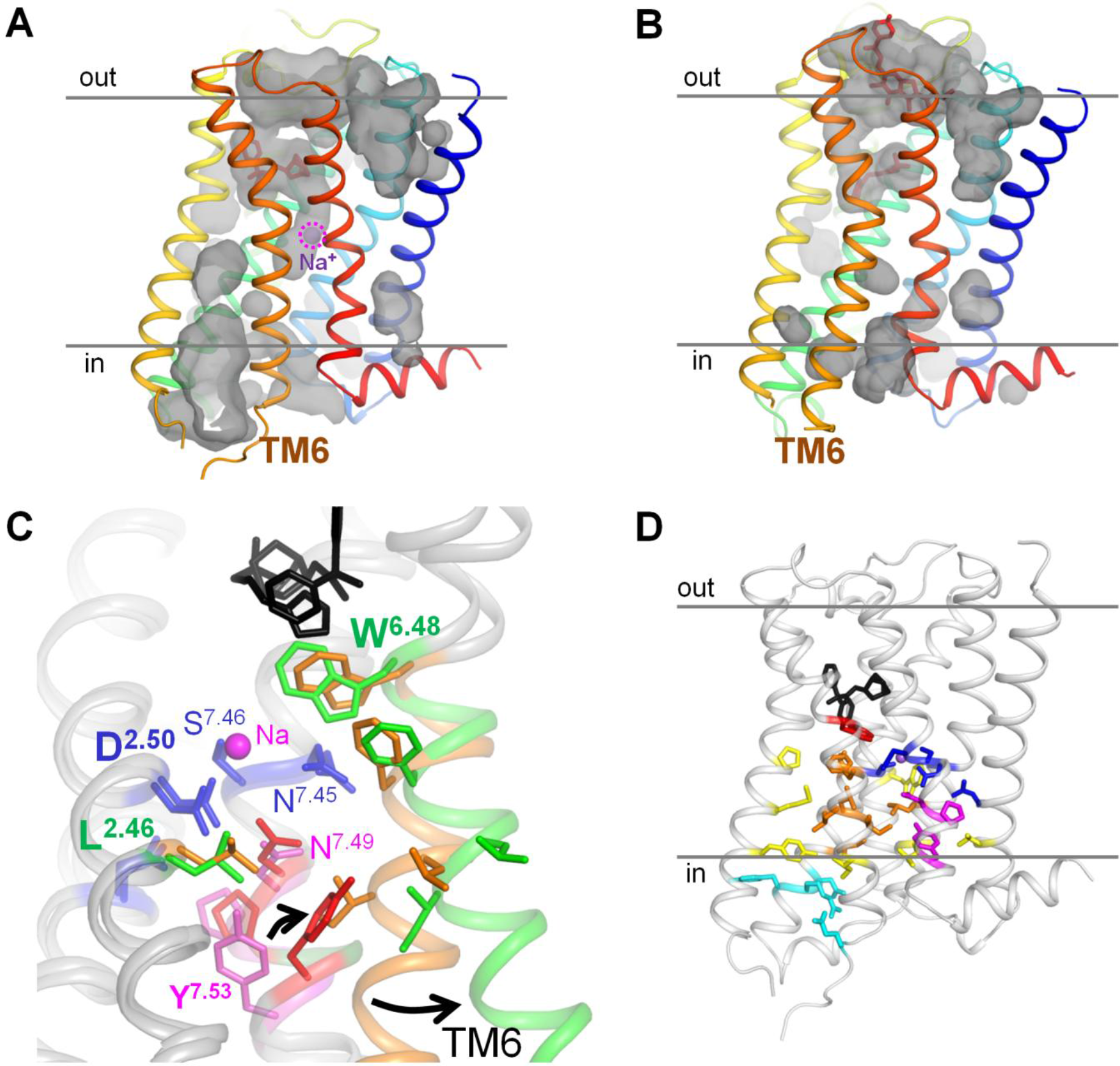
Active and inactive conformations of GPCRs. **A, B**, conformations of the muscarinic acetylcholine receptor M_2_ in the in the inactive state (A, PDB 3UON [14]) and active state (B, PDB 4MQT [13]) with cavities shown; the lines indicate the boundaries of the membrane hydrophobic layer as taken from OPM database [29], “out” indicates the extracellular side, “in” indicates the cytoplasmic side. **C**. Superposition of the structures of M_2_ receptor in the inactive state (the first layer of hydrophobic residues around the Na^+^-binding site colored ochre and residues of the NPxxY motif colored magenta) and in the active state (the first layer of hydrophobic residues around the Na^+^-binding site colored green and residues of the NPxxY motif colored red). The protein is shown as a gray cartoon, the agonist LY2119620 and antagonist 3-quinuclidinyl-benzilate are in black, Na^+^-coordinating residues are colored dark blue. **D**, Overall location of the conserved motifs and hydrophobic residue clusters in GPCRs, shown on the structure of M_2_ receptor in the inactive state (PDB 3UON). The Trp6.48 residue is colored red, the second layer of hydrophobic residues is colored yellow, the residues of the DRY motif and the ionic lock are in light blue. Otherwise the color code as on panel C.

The Na^+^ binding residues are highly conserved among class A GPCRs [26, 28], indicating that the Na^+^ ion must be functionally important. Indeed, replacement of the key Na^+^-binding Asp2.50 residue in helix 2 (see Fig. 1) facilitates binding of agonists by increasing the association constants by 2-3 orders of magnitude in many, albeit not all, cases [26, 28, 30–32]. In many cases, Na^+^ depletion increased the basal activity of the respective receptor in the absence of an agonist, suggesting that the Na^+^ ion stabilizes the inactive state of the receptor [26, 30, 32, 33]. This interpretation is consistent with the structural data that show that the Na^+^ ion is not observed in GPCRs in an active conformation [13, 26, 28, 34, 35].

Several years ago, Katritch and colleagues made a seminal suggestion that the Na^+^ ion does not return into the extracellular medium but instead gets released into the cytoplasm upon GPCR activation [26]. The suggested mechanism, however, would require a transient opening of a conduit for the Na^+^ ion. Although the available structures of activated, Na^+^-lacking GPCRs [13, 34, 35] show no such a conduit, several computer simulations indicated a possibility of a transient water-filled channel connecting the Na^+^-binding site with the cytoplasm [24, 36, 37]. The proposed ability for the Na^+^ ion to traverse the GPCR molecule is also supported by the structural similarity between GPCRs and Na^+^-translocating bacterial rhodopsins (Fig. S1), which suggests their common origin [7] and a common ability to translocate the Na^+^ ion.

Since the cytoplasm is negatively charged relatively to the extracellular medium, transfer of the Na^+^ cation into the cell could give an energy boost for the GPCR activation [26]. Depolarization of the membrane would thus prevent the GPCR activation. This kind of behavior has been, indeed, observed with many GPCRs. Still, some GPCRs, by contrast, were activated by depolarization, whereas some others were insensitive to it; see [38, 39] for reviews. Hence, the relation between the transmembrane gradient of sodium ions and GPCR activation deserves further clarification.

Unfortunately, the anticipated translocation of only one Na^+^ ion per activation event translates into a very weak electric current, which hampers the experimental tracking of this process. In the absence of direct experimental data on the Na^+^ ion translocation, we have addressed it through modeling and comparative structural analyses.

Building on their similarity with Na^+^-translocating microbial rhodopsins, we modeled class A GPCRs as facultative sodium carriers. The model implies that the bound Na^+^ ion, upon activation, can either slip in into the cytoplasm via a transiently opened passage (the carrier-on mode) or return to the extracellular side (the carrier-off mode). Just by varying the dissociation constants for the agonist and the sodium ion, our model quantitatively describes the available data on both activation and suppression of GPCRs by membrane voltage. In addition, by combining evolutionary analyses with structural comparisons of GPCRs, we have identified the strictly conserved leucine residue in the second transmembrane helix (Leu2.46 in class A GPCRs) as the key player in coupling sodium translocation to receptor activation.

In summary, this study proposes a mechanism that links GPCR activation, via Na^+^ translocation, with the energy of membrane potential thereby offering an explanation for the high sensitivity and selectivity of class A GPCRs.

## Results

### 1. Modeling Na^+^ translocation in GPCRs

We developed a model of GPCR activation that is analogous to earlier approaches to modeling energy-converting enzymes [40] and GPCRs [41–44] but takes into account the possibility of electrogenic translocation of a single Na^+^ ion by a GPCR concomitantly with its activation.

The model considers a GPCR ensemble large enough for thermodynamic modeling. Each receptor exists in a steady state balance between its active and inactive states; agonists shift the distribution towards the active state, whereas Na^+^ binding stabilizes the inactive state (Fig. 2A). Both the inactive and active states were shown to exist as series of fast exchanging conformation sub-states [2, 45–47]; for simplicity, we do not consider these sub-states in the model. Also for simplicity it is just assumed that an activated GPCR triggers the signaling cascade; the interaction of the active state with any other component (G-protein or arrestin) is not modeled. The model includes three binary transitions: (i) activation of the receptor, (ii) binding of the Na^+^ ion, and (iii) binding of a signaling molecule (e.g. an agonist). These three transitions can be presented as a cubic graph with 8 separate states (Fig. 2B). Each of the transitions is determined by its equilibrium constant: the receptor activation constant L, the Na^+^ association constant M, and the agonist association constant N (Table 1).

As shown in Fig. 2A, a GPCR has two operation modes: in the carrier-on mode (mode 1), the Na^+^-binding site can communicate with the cytoplasm, whereas in the carrier-off mode (mode 2), the Na^+^-binding site communicates only with the extracellular side. If the receptor operates in the carrier-off mode, all its eight states are in a thermodynamic equilibrium, and we can apply to them the principle of detailed balance (the forward and backward rates of transitions match each other). If the GPCR can translocate a Na^+^ ion across the membrane (carrier-on mode), then, under conditions of a nonzero transmembrane electrochemical sodium potential difference (sodium gradient), the slowest transitions in Fig. 2B are not in equilibrium (the forward and backward rates of these transitions do not match each other).

Our model assumes that the rate-limiting step is the slow "activating" conformational transition from the R state to the final R* state [44]. The ligand binding to the initial encounter state R, binding/release of the Na^+^ ion, and conformational transitions within inactive and active states are considered rapid as compared to the R to R* transition, which appears to be coupled with the pivotal movement of helix 6 in most class A GPCRs [2, 9, 19–25]. In the case of opsin, the pivotal movement occurred slower than at 10^−4^ s with the activation energy as high as 60 kJ/mol [20, 48]. The slowness of the transition was also confirmed by NMR data [22, 46]. The assumption of the slow activating transition is in agreement with both the induced-fit model [43, 44, 47] and the conformational selection model [46]; these two models appear to describe the operation of most GPCRs. Then, at equilibrium, receptor activation is determined by the changes in the free energy of particular states and can be described using the principle of detailed balance, according to which each elementary process should be equilibrated by its reverse process at equilibrium. Since activation of the receptor is much slower than any other transitions in the model, the detailed balance principle can be applied separately to all inactive and all active receptor states (the left-hand and right-hand sides of the cubic diagram in Fig. 2B, respectively).

**Figure 2.**
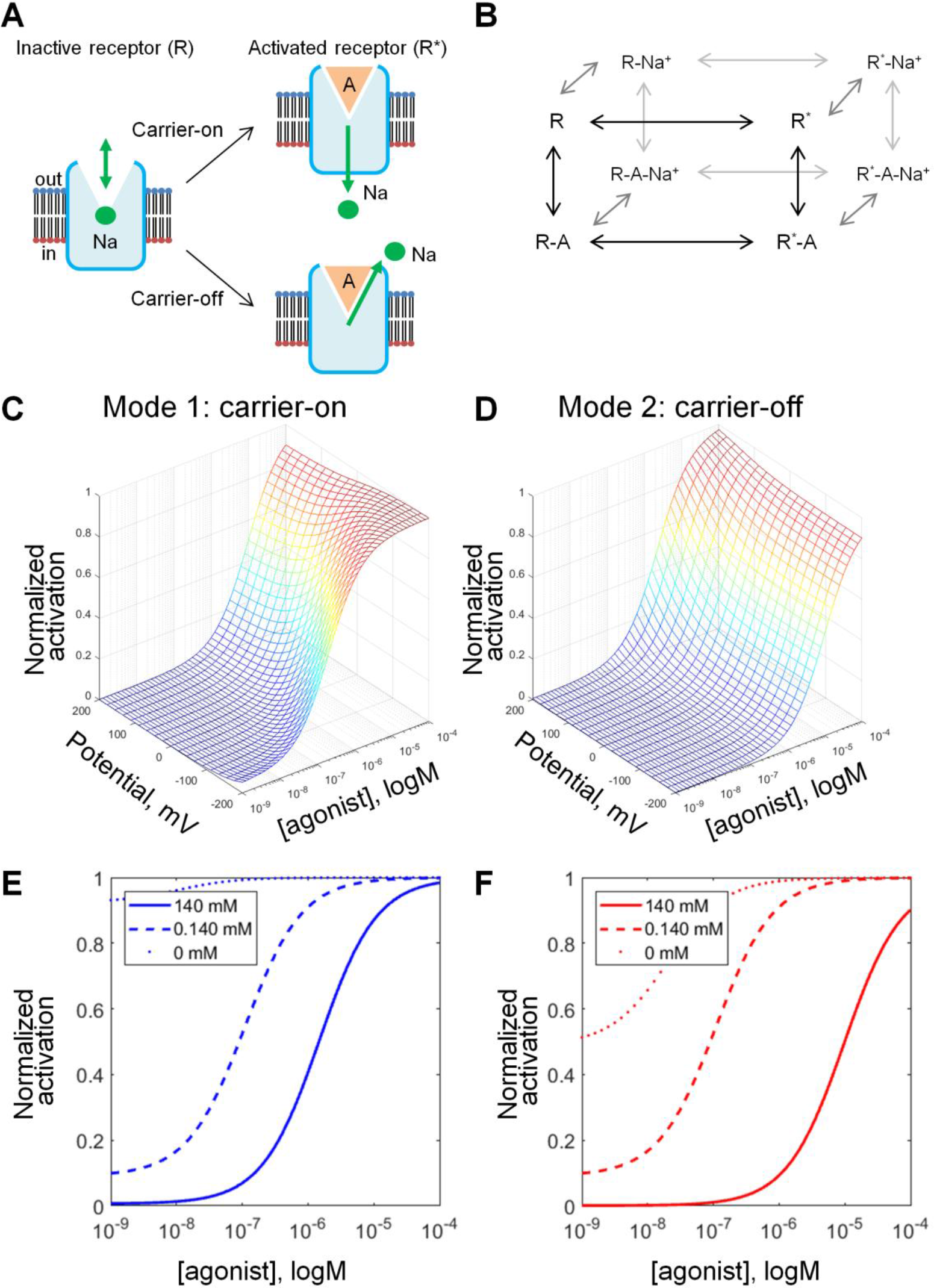
Kinetic models describing the behavior of the Na^+^ ion upon GPCR activation. **A.** Schematic representation of GPCR activation in carrier-on and carrier-off modes. In both modes, the binding of a Na^+^ ion by a GPCR is much more likely in the inactive state than in the active state. When activation of the receptor is triggered by the agonist binding, Na^+^ leaves its binding site. In mode 1 (carrier-on), Na^+^ escapes to the cytoplasm, thus penetrating the membrane. In mode 2 (carrier-off), Na^+^ escapes back into the extracellular space. **B.** General scheme of the model that describes the effect of the agonist and Na^+^ as allosteric modulators on the distribution of the receptor between the active (R*) and inactive (R) states. **C, D**. Effect of the membrane potential on the concentration-response curves of GPCR activation as calculated for the mode 1 (carrier-on) (C) and mode 2 (carrier-off) (D). The curves were plotted with constant allosteric coefficients (α,β,γ,δ) from Table 1, M=10^5^, N=10^5^, and θ=0.65; **E, F**, Effect of [Na^+^]_out_ on the concentration-response curves of GPCR activation, membrane voltage was set to −90 mV, other parameters as in Table 1.

In the absence of a ligand, the fraction of receptors in the R* state is determined by the equilibrium constant L. Generally, L should be about 1, otherwise the receptor will be blocked in one of the two states. The exact value of L, however, is unknown; its determination requires experimental data on the distribution of active and inactive states in the absence of both Na^+^ ions and ligands, which are difficult to obtain. In this study, we set the value L=1. This value was chosen to produce very little activity at low concentrations of the agonist in the presence of Na^+^ ions, but also to exhibit some activity in their absence, since this would most closely mimic the behavior of experimental systems [26, 27, 30, 32, 33]. Specifically, β_2_-adrenergic receptors were biased towards inactive conformation both when studied by ^19^F-NMR and double electron-electron resonance spectroscopy (detergent-stabilized samples, 100 mM NaCl) [2] and by single-molecule monitoring of fluorescent probes (nanodisc-embedded receptors, 150 mM NaCl) [45].

The initial values of the agonist and sodium association constants were set to N=10^7^ M^−1^ and M=10^3^ M^−1^, respectively, in accordance with the physiologically relevant agonist and sodium concentrations, see [27, 49, 50].

The allosteric coefficients α, β, γ, and δ describe the extent of coupling between the transitions, i.e. their interdependence, see Table 1. The coefficient α reflects the effect of Na^+^ ion binding on the receptor activation. Based on experimental evidence, Na^+^ ions inhibit receptor activation [26, 28, 30–32], thus this coefficient should be ≪ 1. The coefficient β reflects the effect of agonist binding on the activation of the receptor. As agonist binding stimulates receptor activation, this value was expected to be ≫ 1. The coefficient γ reflects the effect of agonist binding on the Na^+^ ion binding. Available crystal structures of GPCR with Na^+^ ion bound usually do not contain an agonist molecule, whereas the structures of the same proteins with agonist bound have no space for a Na^+^ ion [26]; therefore, the value of γ was expected to be ≪ 1. The triple allosteric interaction coefficient δ reflects the coupling between all three processes. Probabilities of individual states are also affected by Na^+^ concentrations, which were set at physiological values of [Na]_out_= 140 mM and [Na]_in_=10 mM (Table 1). Further details of the model and its solution are described in **Methods**.

**Table 1.** Initial parameters of the model

| Parameter | Value | Description<sup>a</sup> |
| --- | --- | --- |
| M | $10^3 \text{ M}^{-1}$ | Association constant of the sodium ion (initial value) |
| N | $10^7 \text{ M}^{-1}$ | Association constant of the agonist (initial value) |
| L | 1 | Receptor activation constant |
| $\alpha$ | $10^{-3}$ | Intrinsic efficacy of sodium: ratio of affinity of sodium for $R^*$ and R |
| $\beta$ | $10^3$ | Intrinsic efficacy of the agonist: ratio of affinity of the agonist for $R^*$ and R |
| $\gamma$ | $10^{-2}$ | Binding cooperativity between the sodium ion and the agonist: ratio of affinity of A for R-Na and R, or of Na for R-A and R |
| $\delta$ | $10^2$ | Activation cooperativity between the sodium ion and the agonist: ratio of affinity of A for $R^*$-Na and R-Na, or of Na for $R^*$-A and R-A |
| $\text{Na}^+_{\text{out}}$ | 140 mM | $[\text{Na}^+]$ in the extracellular medium |
| $\text{Na}^+_{\text{in}}$ | 10 mM | $[\text{Na}^+]$ in the cell cytoplasm |

<sup>a</sup> The names of parameters are taken from ref. [42].

Fig. S2 shows how the shape of activation curves varies with variation of parameters α, β, γ, and δ in the carrier-on mode. These concentration - response curves report the sensitivity of the receptor: they show how much agonist is needed to activate the receptor. Fig. S2 shows that increasing the value of δ, which characterizes the coupling between the (i) binding of agonist, (ii) release of the Na^+^ ion into the cytoplasm and (iii) activation of the receptor, increases the sensitivity of the receptor (less agonist is needed for activation, see Fig. S2D).

Based on comparison of the fit curves (Fig. S2) with experimentally measured typical activation curves for several GPCRs, we set the initial values of α, β, and γ to 1000, 0.001, and 0.01, respectively (Table 1). Then the product αβγ equals 0.01, which means that the concurrent binding of the Na^+^ ion and agonist shifts the receptor into the inactive state. Such an inactivation could be avoided by setting δ, the triple allosteric interaction coefficient that reflects the coupling between all three processes, at ≫ 1. The value of δ was initially set to 100, so that the product of multiplication of all coefficients was 1, see Methods for details.

Because of the charge of the Na^+^ ion, the probabilities of states in Fig. 2B would also be affected by membrane voltage. Since cell cytoplasm is charged negatively relative to the extracellular medium, membrane voltage pushes the Na^+^ ion from the extracellular side to the cytoplasmic side of the membrane. Hence, membrane voltage favors the translocation of the Na^+^ ion (i) from the outside into the Na^+^-binding site and (ii) from the Na^+^-binding site into the cytoplasm. In contrast, the retreat of the Na^+^ ion to the extracellular site would be hampered by membrane voltage. The energy gap imposed by membrane voltage can be then defined as:

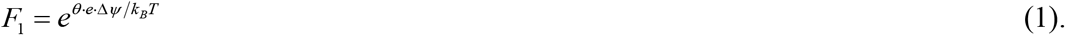

where Δψ is the membrane voltage, *e* is the cation charge, and the coefficient θ reflects the depth of the Na^+^-binding site in the membrane, with θ = 0 corresponding to the Na^+^ ion on the extracellular side of the membrane and θ = 1 corresponding to the Na^+^ ion on the cytoplasmic side of the membrane.

Based on the actual position of the Na^+^ ion in available crystal structures (Fig. 1) and assuming a symmetrical distribution of dielectric permittivity value along the transmembrane axis, we have set θ=0.65 for the Na^+^-binding site in class A GPCRs.

Hence, in the inactive conformation, the membrane voltage promotes Na^+^ binding from the extracellular medium by pushing the cation inside the membrane with the force F_1_ (see Eq. 1). In the carrier-on mode, membrane voltage could push the Na^+^ ion from its binding site in the middle of the membrane into the cytoplasm with the force of

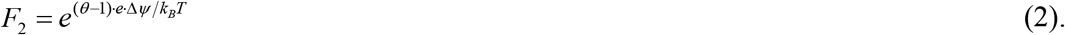

As shown in Fig. 2C and 2B, the same initial parameters from Table 1 yield different activation curves and different dependence on the membrane voltage depending on the operation mode. In the carrier-on mode, the receptor displays the highest sensitivity at the membrane voltage of −200 mV, while neutral and positive voltage values lead to weaker activation, with more agonist required to reach the same activation level. In the carrier-off mode, the dependence of activation on the membrane voltage is opposite: the receptor is more efficient at positive voltage values; here the difference in activation curves at different voltage values is much less pronounced than in the carrier-on mode.

The curves in Fig. 2C, D and Fig. S2 were calculated at fixed physiological concentrations of Na^+^ ions in the external medium and in the cytoplasm (Table 1). In Fig. 2, panels E and F show the activation curves as a function of external Na^+^ concentration. Decreasing the Na^+^ levels increases both the sensitivity of the receptor and the probability of receptor activation in the absence of agonists, in agreement with experimental observations [26, 28, 30–32].

#### Comparison with experimental data

Several studies have reported voltage-dependent signaling in different GPCRs, see Table S1 and [38, 39] for reviews. The membrane voltage of physiological sideness (with cytoplasmic side negatively charged) usually activated GPCRs, although in some cases, e.g. in the muscarinic acetylcholine receptor M_1_, membrane voltage decreased the activity of the receptor [49, 51, 52]. In early studies of voltage-dependent behavior of GPCRs, their activation was followed via downstream reactions, e.g. by measuring the conductance of the GPCR-regulated ion channels [38, 39]. However, the potential voltage-dependence of the channels themselves could make the data interpretation ambiguous. Therefore we limited our scope to the data on voltage dependence of GPCR activation as obtained by using FRET-based biosensors that responded to the outward movement of the transmembrane helix 6 which is directly linked to receptor activation [34, 49, 50]. Using this technique, Rinne and colleagues showed that membrane voltage increased the sensitivity of the α_2A_ adrenoreceptor to norepinephrine [50]. In addition, membrane voltage increased the sensitivity of muscarinic acetylcholine receptors M_3_ and M_5_ to their full agonists acetylcholine and carbachol, but decreased the sensitivity to the same full agonists in the case of the muscarinic acetylcholine receptor M_1_ [49].

We tested whether our model could describe the data of Rinne and colleagues [49, 50]. To perform the fitting, experimental data points were extracted from respective publications. In each case, fits were separately performed for the carrier-on and carrier-off modes, respectively (Fig. 2). For each mode, two sets of data points (as measured at two voltage values, e.g. −90 and +60 mV) were fitted. The two fit curves were calculated with the same parameter sets (see Table 1), with the only difference being the voltage values that were taken from respective experimental data. Both curves were fitted simultaneously by varying only two parameters, namely the Na^+^ binding constant M and agonist binding constant N, which were kept the same for both voltage values in each case. All other parameters were as in Table 1. In addition, we performed separate fits with β=100, see Table 2.

The fit curves are shown in Fig. 3 and the fit parameters are listed in Table 2. In each case, one of the two tested modes provided a good fit. In cases when the membrane voltage increased the sensitivity of the receptor, the experimental data could be fit by the model in carrier-on mode. For the data in Fig. 3C, where the sensitivity was decreased by membrane voltage, the carrier-off mode provided a good fit. Varying the θ value, which was initially determined from structure analysis, did not lead to notable fit improvements (data not shown). It is noteworthy that the agonist association constants in Table 2 correspond to the so-called intrinsic association constants (K_a_), which could be much smaller than the observable association constants (K_obs_), see [44, 53].

**Figure 3.**
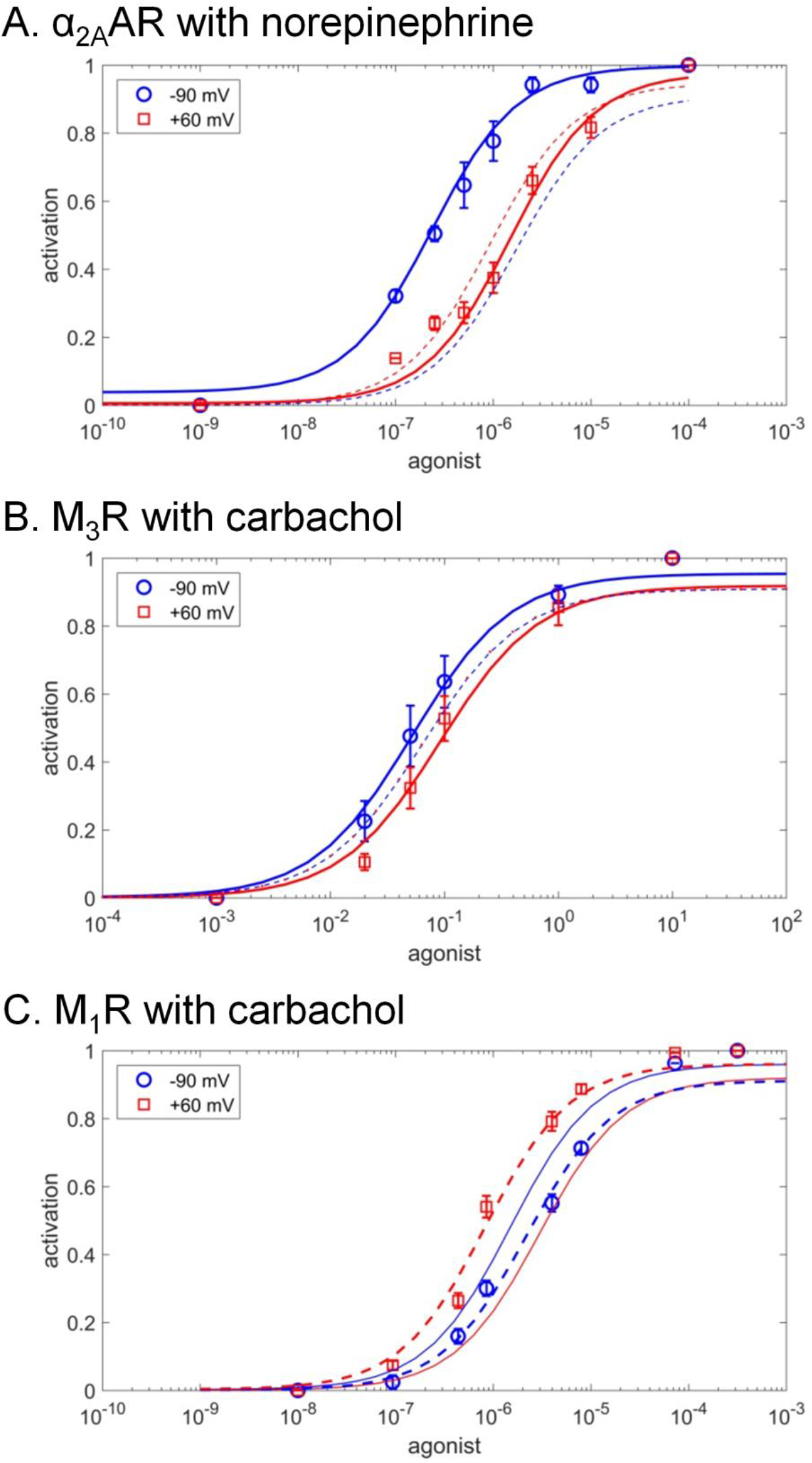
Fitting experimental data of voltage-sensitive GPCR activation. The fits for the carrier-on mode 1 are shown by solid lines, the fits for the carrier-off mode 2 are shown by dashed lines. **A.** α_2A_ adrenoreceptor activation by full endogenous agonist norepinephrine, experimental data from [50]. **B.** Muscarinic M_3_ receptor activation by full agonist carbachol experimental data from [49]. **C.** Muscarinic M_1_ receptor activation by full agonist carbachol, experimental data from [49].

**Table 2.** Sodium ion and agonist binding constants as obtained from fitting experimental data

| Receptor – agonist pairing | Mode | $\beta=1000$ | | $\beta=100$ | |
| --- | --- | --- | --- | --- | --- |
|  |  | Na<sup>+</sup> association constant, M<sup>-1</sup> | Agonist association constant, M<sup>-1</sup> | Na<sup>+</sup> association constant, M<sup>-1</sup> | Agonist association constant, M<sup>-1</sup> |
| $\alpha_{2A}$-AR with norepinephrine | 1, carrier-on | $2.4 \cdot 10^4$ | $2.6 \cdot 10^5$ | $8.7 \cdot 10^3$ | $1.1 \cdot 10^6$ |
| M<sub>3</sub> muscarinic receptor with carbachol | 1, carrier-on | $3.5 \cdot 10^5$ | 8.4 | $3.1 \cdot 10^5$ | 87.9 |
| M<sub>1</sub> muscarinic receptor with carbachol | 2, carrier-off | $2.1 \cdot 10^7$ | $6.4 \cdot 10^5$ | $2.2 \cdot 10^5$ | $2.6 \cdot 10^6$ |

By using essentially the same set of parameters from Table 1 and varying only the values M and N, it was possible to quantitatively fit also the previously obtained experimental data on the influence of membrane voltage on the activation of GPCRs as traced via downstream reactions (Table S1). An example of such a fit is presented in Fig. S3.

Our modeling of experimental data speaks against the obligatory coupling between the activation of a GPCR and transmembrane translocation of a Na^+^ ion into the cytoplasm. It appears that in some GPCRs, the Na^+^ ion can retract, upon activation, into the extracellular medium. In most studied cases, however, the experimental data are better described by the carrier-on operation mode where binding of an agonist is mechanistically coupled to the voltage-driven translocation of the Na^+^ ion into the cytoplasm.

## Discussion

### 1. Modeling the behavior of GPCRs

Here, staying in the traditional bioenergetics framework, we considered Na^+^-dependent GPCRs as potential electrogenic Na^+^ carriers. The ability of GPCRs to provide a passage for a Na^+^ ion follows from (i) the results of molecular dynamics simulations [24, 36, 37], (i) the evolutionary relatedness of GPCRs to, and structural similarity with, such dedicated transporters as Na^+^-dependent bacterial rhodopsin (NR) and channelrhodopsin [7], see the discussion below and Fig. S1, S5; and (iii) the data on voltage-driven charge displacements within GPCRs [52, 54]. Vichery and colleagues argued that the so-called “gating currents” or “sensory currents”, measured in response to imposed membrane voltage with some GPCRs [52, 54], may reflect the movement of a Na^+^ ion from its binding site into the cytoplasm [39].

To clarify the benefits from Na^+^ translocation, let us consider the thermodynamics of a membrane receptor. An "ultimately sensitive" receptor should stay inactive in the absence of agonists and get activated (e.g. by changing its conformation from R to R*) in response to the very first arriving molecule(s) of agonist. In such a system, the free energy needed to drive the conformational change is provided by the binding of agonist; the amount of this free energy is, however, small when the concentration of agonist is comparable with its dissociation constant. Indeed, the free energy of binding can be determined as

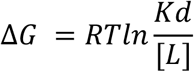

where *Kd* is the dissociation constant of the ligand (e.g. agonist) and [*L*] is the concentration of the ligand. Then, if the dissociation constant of an agonist is 10^−7^ M, the free energy of its binding will be zero at 10^−7^ M of agonist and only −6 kJ/mol at 10^−6^ M of agonist. With such a small energy input, the activation would be possible only if the equilibrium constant between R and R* is small (i.e. close to unity, as in our modeling). Then, however, because the energy of thermal fluctuations is 2.5 kJ/mol (one kT) at room temperature, 10% of receptors would stay constantly activated even in the absence of agonist and produce spurious noisy signal.

Selective stabilization of receptors in their inactive state by Na^+^ ions, which are abundant outside the cell, would help to silence the intrinsically noisy GPCRs in the absence of agonist molecules. However, such a noise reduction, at the same time, would decrease the sensitivity of GPCRs. To activate a Na^+^-blocked receptor, proportionally higher levels of agonists would be needed. This conundrum can be solved only by invoking an external source of free energy and coupling it to the receptor activation. The data on voltage dependence of some GPCRs in Fig. 3A,B and S3 indicate that these GPCRs use the energy of transmembrane electric field to increase their sensitivity. At physiological membrane voltage of about −100 mV, the strength of the electric field that pushes the Na^+^ ion across the membrane is approx. 10^7^ V/m. The energy of this field could be, however, used only if the receptor activation is mechanistically coupled to the displacement of the ion.

How much free energy could be gained this way *in vivo*? In experiments where the voltage was varied by 150 mV (between −90 mV and 60 mV, see Fig. 3, S3), only the membrane voltage, but not the Na^+^ concentrations, were varied. *In vivo*, the movement of the Na^+^ ion would be driven both by the voltage on the cellular membrane Δψ (~−100 mV) and concentration difference of Na^+^ (140 mM outside *versus* < 10 mM inside). By analogy with the proton-motive force introduced by Mitchell for describing the proton-motive energy conversion [55], the corresponding sodium-motive force (*smf*), in the case of Na^+^-translocating membrane enzymes, could be defined as

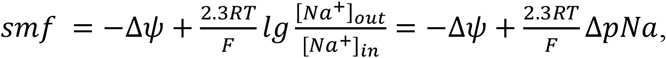

where 2.3RT/F is 59.1 mV at 298°K [56].

In case of GPCRs, the value of *smf* would depend on the extent of the coupled transmembrane charge transfer. After the escape of the Na^+^ ion into the cytoplasm, the Asp2.50 residue is unlikely to stay deprotonated and negatively charged in the middle of the membrane, it would rather accept a proton from the external medium [37]. The corresponding displacement of a proton would be driven by membrane voltage; it, however, could be either coupled to the activation of the receptor or not. If proton displacement uncoupled (e.g. because protonation of Asp2.50 is slow and happens after the activation of the GPCR), the amount of free energy derived from the Na^+^ translocation could be estimated as (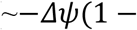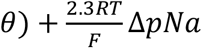) or ~ 120 meV. As it follows from Fig. 2C, the *smf* of 120 meV, at δ ≫ 1.0, would increase the receptor sensitivity up to an order of magnitude. If the reprotonation of Asp2.50 is coupled to the activation, as argued by Vickery and colleagues [37], the respective *smf* of ~ 170 mV 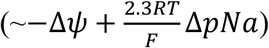 could increase the sensitivity by almost two orders of magnitude (see Fig. 2C). Furthermore, reprotonation of Asp2.50 could be productive in some GPCRs and futile in other GPCRs contributing to the variations of voltage effects, as reviewed in [37, 39].

Our modeling of experimental data (Fig. 3 and S3) supports the idea that Na^+^-translocating GPCRs, when operating in the carrier-on mode, could use the energy of the transmembrane electric field to amplify the signal. Devices that use the energy of external electric field to amplify a weak signal are called field effect transistors.

Additionally, the same electric field, by preventing the escape of the bound Na^+^ ion to the extracellular side and stabilizing the inactive conformation, would decrease the noise in the absence of agonist. Hence, Na^+^-dependent GPCRs can work as electrochemical field effect transistors in which the electric field is additionally used to suppress the noise.

There are even more benefits from coupling translocation of the Na^+^ ion with receptor activation. Since the equilibrium constant between the active and inactive states of GPCRs is low (Table 1), the equilibrium, in principle, could be shifted towards the active state in multiple ways in response to binding of various ligands to different patches in the ligand-binding pocket. The above-described amplification mechanism, however, would be involved only when the binding of an agonist molecule would prevent the retreat of the Na^+^ ion into the extracellular media upon activation [26]. The Na^+^ binding site is connected with the extracellular space by a polar cavity (Fig. 1A, B), so that the retreat of a Na^+^ ion through this cavity upon activation is mechanistically easier than its corkscrewing in the opposite direction through the layer of hydrophobic residues - unless the agonist binds in such a specific way that the retreat of the Na^+^ ion is blocked and it is forced to escape in the opposite direction. Apparently, only some specific modes of agonist binding would prevent the retreat of the Na^+^ ion and thus enable the coupling between its electrogenic translocation into the cell and the receptor activation. The signal of such agonist molecules would be amplified by electric field (carrier-on mode), whereas the signal of other, e.g. partial agonists would be weakened by the field (carrier-off mode), which would dramatically increase the chemical selectivity of the receptor.

It could be anticipated that the endogenous agonists of studied GPCRs should be among those effectors whose signals are amplified by membrane voltage. Indeed, in the muscarinic acetylcholine receptor M_2_, the membrane voltage potentiated the signal from the endogenous full agonist acetylcholine, but decreased the signal from the drug pilocarpine, a partial agonist, see Table S1, Fig. S3 and [57, 58]. Additionally, the voltage sensitivity of the M_2_ receptor was shown to be altered by mutations in the orthosteric ligand-binding site, indicating a direct connection between the agonist binding and voltage effects [57].

In the case of the muscarinic acetylcholine receptor M_3_, the membrane voltage increased the sensitivity to full agonists acetylcholine (endogenous) and carbachol (artificial), but decreased the sensitivity to choline or pilocarpine, see Table S1, Fig. 3B and [49]. In support of their suggestion that voltage-sensitivity is defined by the specific binding mode of each signaling molecule to the receptor, Rinne and coworkers have shown that the replacement of Asn6.52 in the M_3_ receptor by Gln reversed the voltage effect in case of the artificial agonist carbachol but did not affect the behavior of the endogenous agonist acetylcholine [49]. The authors concluded that their data buttress the importance of the 6^th^ helix for the mechanics of acetylcholine receptors.

Similarly, membrane voltage increased the sensitivity of dopamine D_2S_ receptor to its native agonist dopamine, but decreased the sensitivity to β-phenethylamine, *p*- and *m*-tyramine, see Table S1 and [59–61].

By contrast, activation of the muscarinic acetylcholine receptor M_1_ both by endogenous agonist acetylcholine and artificial agonist carbachol was depressed by membrane voltage, see Fig. 3C and Table S1. This behavior could be better described by the carrier-off mode, where the Na^+^ ion retreats into the extracellular medium upon activation. It appears that the M_1_ receptor does not translocate Na^+^ even in the presence of its endogenous agonist, which may have sense if the activity of this receptor is suppressed by membrane voltage.

Finally, Table S1 also reports several cases where receptors were insensitive to membrane voltage. Mechanistically, this could happen if the displacement of the Na^+^ ion - in either direction - is uncoupled from the activation (δ≪1.0). For instance, Na^+^ ion could retreat before the major conformational change/activation occurs.

In summary, the suggested model allowed us to qualitatively describe the data on voltage dependence of several Na^+^-dependent GPCRs by using the same set of fixed parameters for receptor activation and by varying only the affinities to the agonist and the Na^+^ ion. It appears that the translocation of a Na^+^ ion across the membrane and into the cytoplasm concomitantly with the receptor activation increases both the sensitivity and selectivity of GPCRs.

### 2. Structural elements of the coupling mechanism

Here we argue that the Na^+^ ion serves as the physical moiety that couples the GPCR with the external electrochemical gradient by pushing, as a cannon ball, through the layer of hydrophobic residues and promoting the conformational change in the receptor. To identify the structural aspects of the coupling mechanism, we superposed crystal structures of the muscarinic acetylcholine receptor M_2_ in its inactive state (PDB entry 3UON [14]) and in the active state (PDB: 4MQT [13]), see Fig. 1C. This pair of structures was chosen because of the clear-cut differences between the inactive and active structures [13, 62] and the reported voltage dependence of muscarinic receptors, see Table S1, Fig. S3 and [51, 52, 63]. The results of our analysis, however, should also be applicable to other Class A GPCRs because of the high sequence identity of the regions surrounding the sodium ion among the members of this class [26]. They share, for example, motifs LxxxD in helix 2, DRY in helix 3, WxP in helix 6, and NPxxY in helix 7, see Fig. 1C,D and [64–66]. Fig. 1D shows the positions of these motifs, whereas Table 3 reports the extent of their conservation.

**Table 3.** Conservation of functionally important residues in Class A GPCRs

| Residue in M<sub>2</sub> muscarinic<br>acetylcholine receptor<br>(PDB: 3UON) | Generic<br>number | Most common residue |  | Second most common<br>residue |  | Residue type, % |
| --- | --- | --- | --- | --- | --- | --- |
|  |  | AA | % | AA | % |  |
|  | Na<sup>+</sup> coordination |  |  |  |  |  |
| Asn41 | 1.50 | N | 98 | S | 1 | Polar, 100 |
| Asp69 | 2.50 | D | 92 | N | 3 | Polar, 98 |
| Ser110 | 3.39 | S | 72 | T | 8 | Polar, 83 |
| Trp400 | 6.48 | W | 68 | F | 16 | Aromatic, 87 |
| Asn432 | 7.45 | N | 67 | S | 11 | Polar, 93 |
| Ser433 | 7.46 | S | 64 | C | 13 | Polar, 72 |
|  | WxP motif |  |  |  |  |  |
| Thr399 | 6.47 | C | 71 | S | 10 | Small, 86 |
| Trp400 | 6.48 | W | 68 | F | 16 | Aromatic, 87 |
| Pro402 | 6.50 | P | 99 | N/A | N/A | Helix kink, 99 |
|  | Hydrophobic shell around the Na<sup>+</sup> pocket |  |  |  |  |  |
| Leu65 | 2.46 | L | 90 | M | 4 | Hydrophobic, 99 |
| Val111 | 3.40 | I | 40 | V | 24 | Hydrophobic, 88 |
| Leu114 | 3.43 | L | 73 | I | 10 | Hydrophobic, 98 |
| Ile117 | 3.46 | I | 56 | L | 16 | Hydrophobic, 99 |
| Ile392 | 6.40 | V | 37 | I | 28 | Hydrophobic, 93 |
| Leu393 | 6.41 | V | 41 | L | 20 | Hydrophobic, 91 |
| Phe396 | 6.44 | F | 75 | V | 4 | Hydrophobic, 92 |
|  | Second hydrophobic shell |  |  |  |  |  |
| Val44 | 1.53 | V | 65 | A | 14 | Hydrophobic, 92 |
| Ile62 | 2.43 | L | 36 | I | 35 | Hydrophobic, 97 |
| Trp148 | 4.50 | W | 96 | F | 1 | Hydrophobic, 99 |
| Pro198 | 5.50 | P | 79 | V | 5 | Hydrophobic, 95 |
| Met202 | 5.54 | I | 33 | M | 30 | Hydrophobic, 90 |
| Tyr206 | 5.58 | Y | 72 | S | 5 | Hydrophobic, 86 |
| Ile389 | 6.37 | L | 38 | V | 21 | Hydrophobic, 91 |
|  | NPxxY motif |  |  |  |  |  |
| Asn436 | 7.49 | N | 72 | D | 20 | Polar, 98 |
| Pro437 | 7.50 | P | 94 | A | 2 | Hydrophobic, 98 |
| Tyr440 | 7.53 | Y | 89 | F | 4 | Aromatic,93 |
|  | DRY motif (ionic lock) |  |  |  |  |  |
| Asp120 | 3.49 | D | 64 | E | 21 | Polar, 97 |
| Arg121 | 3.50 | R | 95 | H | 1 | Polar, 98 |
| Tyr122 | 3.51 | Y | 66 | F | 10 | Aromatic, 87 |

The crystal structures of the muscarinic acetylcholine receptor M_2_ in the active and inactive state (Fig. 1A-C) differ in the conformation of helix 6 and, specifically, positions of Trp400 (6.48), Leu65 (2.46) and the residues of the NPxxY motif (Fig. 1C, residue numbers for the PDB entry 4MQT). In the inactive state of the M_2_ receptor, a large cavity above the sodium site connects it with the extracellular space, while the cytoplasmic cavity protracts only up to the residues of the NPxxY motif. As discussed earlier [24, 36, 37, 68, 69], the connection between the two cavities is blocked by a layer of hydrophobic residues. In Fig. 1, this hydrophobic layer contains Leu65 (2.46), Leu114 (3.43), Val111(3.40), Ile117(3.46), Ile392(6.40), Leu393(6.41) and Phe396(6.44) (Fig. 1C,D). These hydrophobic residues in helices 2, 3 and 6 were previously implicated in controlling the activation-related conformational changes [24, 36, 37, 68, 69]. Most of these residues are conserved in class A GPCRs (Table 3). The above-listed residues of the hydrophobic barrier are supported structurally by a second shell of conserved hydrophobic residues: Val44(1.53), Ile62(2.43), Trp148(4.50), Pro198(5.50), Met202(5.54), Tyr206(5.58), and Ile389(6.37). These residues appear to be functionally important: their positions are almost always taken by hydrophobic residues in the class A GPCRs, and in many cases not just the hydrophobic nature, but even the specific residue types are highly conserved (see Table 3).

In Fig. 1, the hydrophobic layer is centered on Leu65 (Leu2.46) of the conserved LxxxD motif. Leucines in a helix generally prefer one of two rotamers [70], and a rotamer search for Leu2.46 reveals two clearly preferred rotamers at this position (Fig. S4A). After reinterpretation of the X-ray data of the PDB entry 3UON by the PDB_REDO team [71], Leu2.46 adopts the conformation shown in Fig. 4A and S4B, which corresponds to the lower of the two preferred rotamers in Fig.S4A. The upper of the two rotamer positions is not available to Leu2.46 in the 3UON structure because the Leu side chain would clash with Asp2.50 and Tyr7.53. Notably, in the active structure of the same receptor (PDB: 4MQT), Leu2.46 is observed in the upper rotamer, see Fig. 1C, S4C. If Leu2.46 is placed in the upper rotamer in the structure of the inactive receptor as envisioned in Fig. 4B and S4D, the two cavities get connected, indicating that water and small ions could freely traverse the GPCR helix bundle. Hence, formation of a passage for a Na^+^ ion requires releasing the steric hindrances by Asp2.50 and Tyr7.53, which prevent rotation of Leu2.46 into the upper rotamer position.

**Figure 4.**
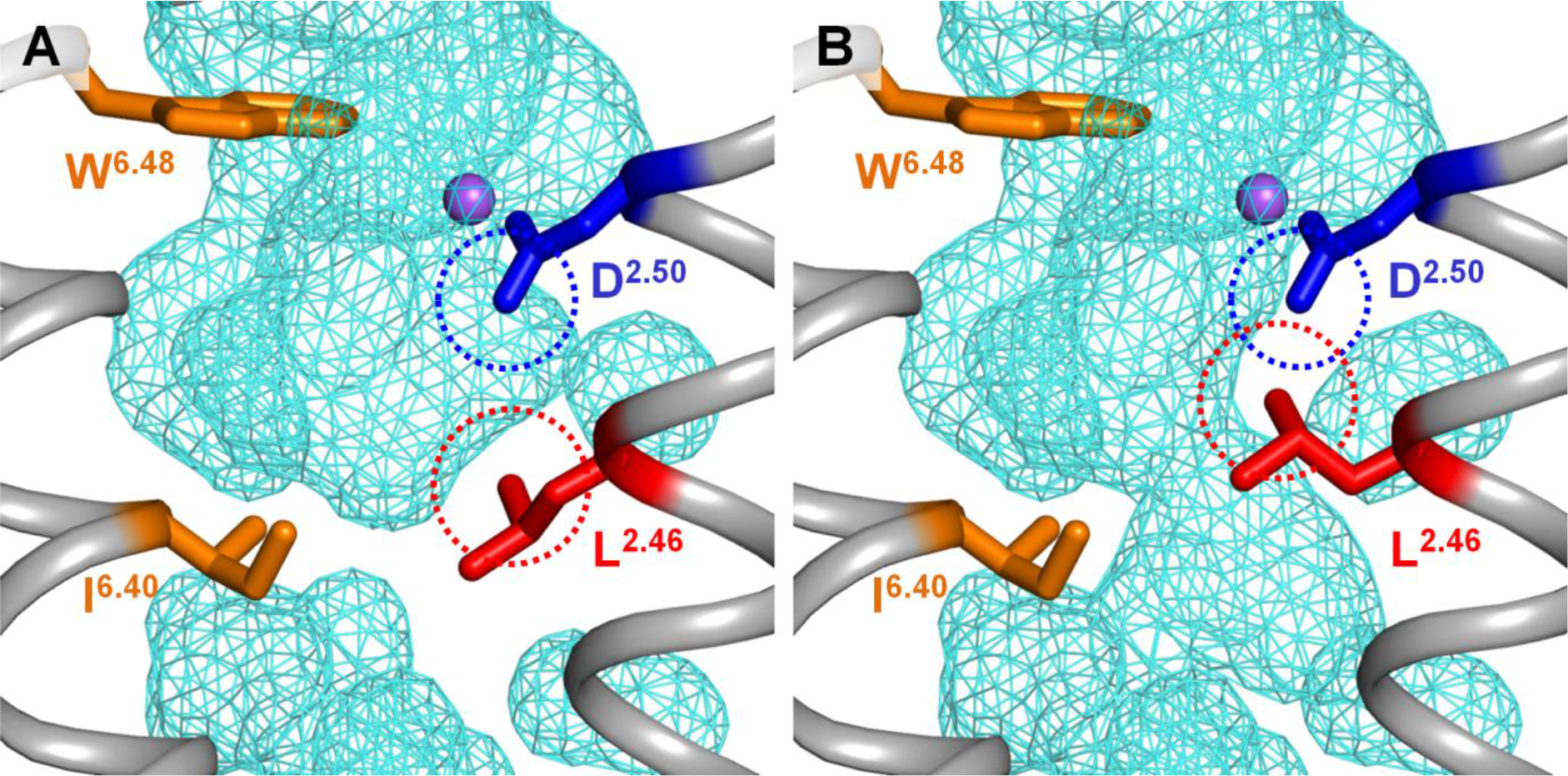
Suggested pathways of Na escape to the cytoplasm upon GPCR activation in the muscarinic acetylcholine receptor M_2_. The protein is shown in gray, Na^+^ ion is in purple, solvent-accessible surface with solvent radius 1.12 Å (corresponding to the Na^+^ ion) is shown as blue mesh. **A**. Sodium-binding cavity in the M_2_ receptor in the antagonist-bound inactive state (PDB 3UON). **B**. A minor rotameric transition of Leu2.46 alone is enough to open the connection between two inner cavities, allowing the Na^+^ ion passage through the layer of conserved hydrophobic residues (see also Fig. S4). The dashed circles mark the Van der Waals radii as calculated by the PyMol v 1.7 software [72].

A minor rotameric transition of the strictly conserved Leu2.46 can open a conduit for voltage-driven translocation of Na^+^ ion into the cell (Fig. 4, S4). Leu2.46 and Asp2.50 form the highly conserved motif LxxxD in helix 2, (Table 3, Fig. 6) where Asp2.50, owing to its negative charge, is the strongest ligand of the Na^+^ ion. While Asp2.50 serves as a Na^+^ ligand, Leu2.46 controls the interface between helices 2 and 3. In the NR, the transmembrane ion passage is formed by helices 3, 6 and 7 [7, 73, 74] and is likely to be contributed by same helices in GPCRs [7, 24, 26]. Based on the structure analysis (Fig. 4, S4), we suggest that the rotameric change of Leu2.46 upon GPCR activation plays a key role in coupling the voltage-driven Na^+^ translocation into the cell with the activation of GPCRs.

Numerous alanine screening experiments, performed on different GPCRs, showed an altered function in Leu2.46 mutants, see e.g. [75]. The results of such experiments, however, are difficult to interpret. If Leu2.46 is indeed, as we suspect, involved in the coupling of GPCR activation with Na^+^ translocation, then its replacement would affect both these processes with an unpredictable outcome for the function. More informative is the observation, obtained upon alanine scanning of the human adenosine A_2A_ receptor, that the mutation of Leu2.46 to Ala (L48A) dramatically increased the thermostability of the receptor [76]. The temperature Tm at which 50% of the solubilized receptor could bind ligand after 30 min thermo-incubation increased from 28.5°C in the wild-type protein to 42.5°C in the Leu2.46Ala mutant. The two second-best mutations increased Tm only to 34.5°C. The L48A mutation apparently fixed the human adenosine A_2A_ receptor in a (thermo)stable, active conformation [76].

Poor stability is a common property of GPCRs, which hindered their crystallization for several decades. This instability is an intrinsic property; the sensitivity of GPCRs is determined by their ability to switch between two different conformations with similar energies. Dramatic stabilization of the active conformation of A_2A_ receptor after replacing Leu2.46 by the smaller Ala indicates that the conformation of the bulky hydrophobic side chain of Leu2.46 is indeed keeping the balance between the active and inactive conformations by serving as an important “weak spot” [77]. At the same time, Leu2.46 appears to control the Na^+^ path (see Fig. 4, S4), which points to this strictly conserved residue as the key coupling moiety in GPCRs.

The coupling between the electrogenic Na^+^ translocation and GPCR activation would be achieved if the agonist binding decreases the affinity of GPCR for the Na^+^ ion. Indeed, keeping the Na^+^ ion in the middle of the hydrophobic membrane is energetically very demanding because of the high desolvation penalty for a positively charged small cation [78]. In the well-studied Na^+^-dependent ATP synthases, binding of the Na^+^ ion in the middle of the membrane requires six ligands [79, 80]; a loss of even one of them transforms a Na^+^-translocating enzyme into a proton-translocating one [79, 81]. The detachment of Na^+^ from its binding site could be mediated by Trp6.48. This residue is strictly conserved in most class A GPCRs (Table 3) and changes its conformation in response to agonist binding, see Fig. 1C and [2, 9, 19–25]. Specifically, in the structure of the δ-opioid receptor (PDB: 4N6H) Trp274(6.48) interacts with the Na^+^ ion via a water molecule [7, 15]; same interaction can be seen in structures of A_2A_ adenosine receptor (PDB 4EIY) and β_1_-adrenoceptor (PDB 5A8E). In all likelihood, Trp6.48 participates in a hydrogen-bonded network also in other class A GPCRs. If the system of bonds around the Na^+^ ion gets destabilized in response to the ligand binding, the further stay of the Na^+^ ion in the middle of the membrane would not be possible. If the retreat path is blocked by the bound agonist, the Na^+^ ion would swing in the pocket until, being pushed by electric field, it enforces the twist of Leu2.46 residue into the upper rotamer. This rotameric transition demands, however, to resolve the steric clash with Asp2.50 that is located just one helix turn away (Fig. 4B). Asp2.50 is engaged in binding of Na^+^ ion and would not get aside as long as the Na^+^ ion stays bound. Hence, the breakage of the bond between the Na^+^ ion and Asp2.50 is a precondition of the isomeric transition of Lys2.46 and opening of the cytoplasmic Na^+^ conduit (carrier-on operation mode, Fig. 2C, 3A, B). Alternatively, the Na^+^ ion could retreat, against the electric field, into the extracellular space. In this case, the electric field would prevent, rather than promote, the GPCR activation (carrier-off mode, Fig. 2D, 3C). After the escape of the Na^+^ ion, Tyr7.53 makes a new hydrogen bond, via a water bridge (with Tyr5.58 in mammals), whereas the Asn7.49 residue of the NPxxY motif turns towards Asp2.50 and forms a new, alternative hydrogen bond network, which is seen in the "active" crystal structures (Fig. 1B, 1C) [22–24]. These new hydrogen bonds prevent the return of the Na^+^ ion and stabilize the active conformation of the receptor, which is accompanied by displacement of several TM helices and closing of the Na^+^-binding pocket and of the cytoplasmic Na^+^ path (Fig. 1B, S4C). As a result, the escape of the Na^+^ ion gets thermodynamically coupled with the GPRC activation.

Our structural inspection of cavities in GPCRs showed that the Na^+^ ion might escape from the helical bundle of GPCR either after reaching the water phase from the cytoplasmic side (dashed line in Fig. 5A) or, even earlier, by slipping between helices 2 and 3 at the level of the DRY motif (dashed line Fig. 5B). In the latter case, the Na^+^ ion would be released into the layer of phosphate groups of phospholipids. Their negative charges, serving as potent alternative Na^+^ ligands, could attract the Na^+^ ion and help it to slide between the helices. For the Na^+^ ion, in fact, there is no need to get any further. Molecular dynamics simulations of phospholipid membranes showed that Na^+^ ions reside among the phosphate groups and compensate their negative charges [82].

We have observed a continuous path for the Na^+^ ion only when the GPCR is in the *inactive* conformation and Leu2.46 is in the upper rotamer typical for the *active* conformation (Fig. 4B, S4D), which explains why the path is not seen either in the active or in inactive structures (Fig. 1, 4A, S4B, C). Owing to the constriction by the second shell of hydrophobic resides (Table 3, Fig. 1D), the probability of a spontaneous opening of the path should be low, which might explain the relatively high activation energy of the conformational change in GPCRs [20, 47, 48]. Otherwise, Na^+^ ions would constantly leak through GPCRs. While the importance of Asp2.50, Trp6.48 and Tyr7.53 for activation of GPCRs and formation of a Na^+^-translocating passage has been widely discussed, see e.g. [2, 22–24, 36, 37], the control function of Leu2.46, to our knowledge, has not been specifically addressed until now.

**Figure 5.**
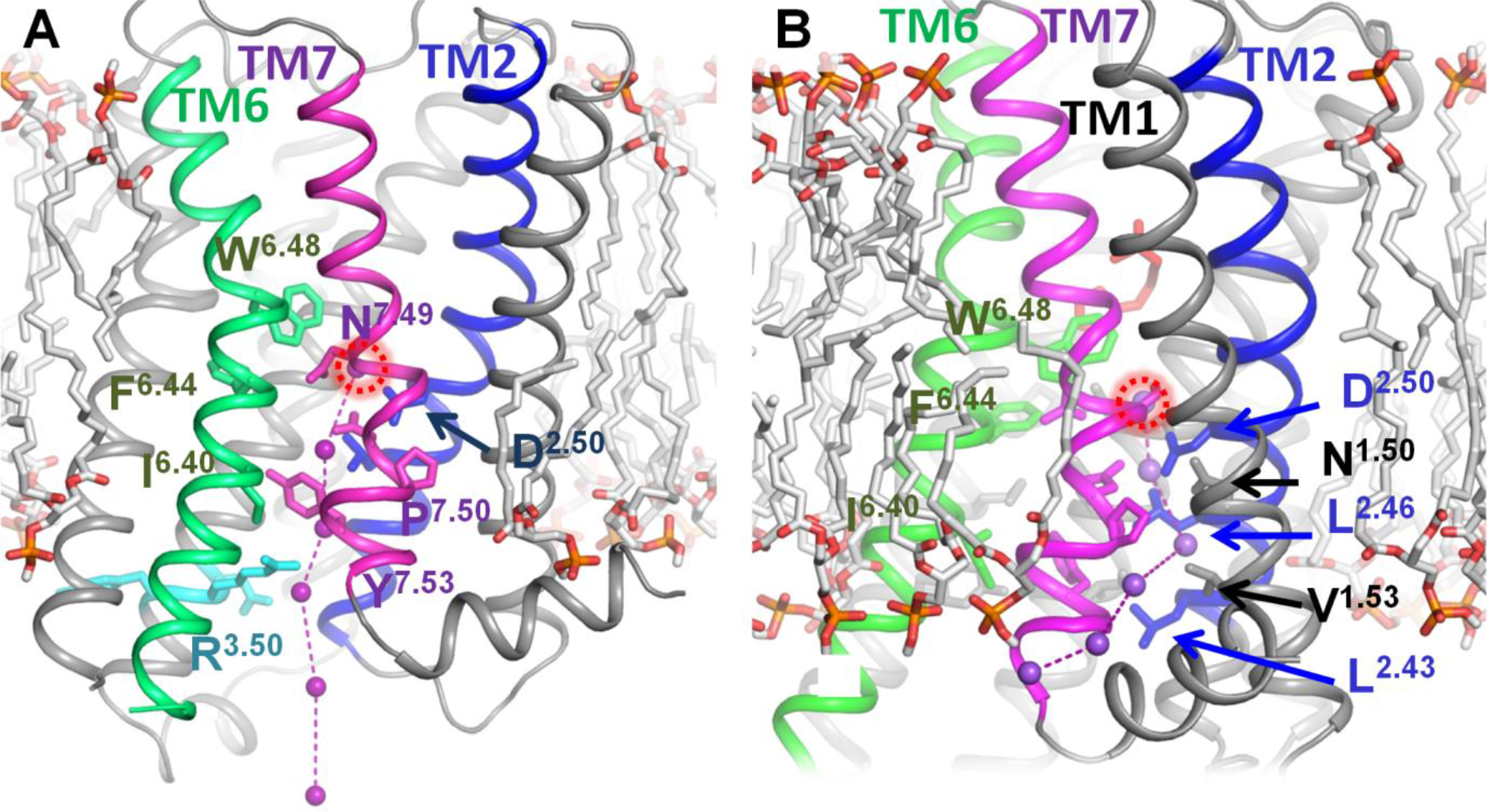
Suggested pathways of Na^+^ escape to the cytoplasm upon GPCR activation, based on the structure of the M_2_ receptor (PDB: 4MQT). **A.**Putative exit pathway for the Na^+^ ion via the center of the heptahelical bundle. **B**. An alternative exit pathway via the pocket between helices 1, 2, and 7. Hypothetical intermediate positions of the Na^+^ ion are shown as purple spheres; the initial position, inferred from the position of Na^+^ in the structures of inactive GPCRs, is indicated by the red dashed circle. Helices 2, 6, and 7 are colored blue, green and purple, respectively; conserved residues listed in Table 3 are shown as sticks. The ionic lock residues are shown in cyan. CHARMM-GUI software [83] was applied to construct the lipid molecules of the membrane surrounding the receptors (shown as grey sticks).

### 3. Evolutionary considerations

Earlier, the structure superposition of the Na^+^-translocating rhodopsin from *Dokdonia eikasta* (NR), and Na-bound GPCRs allowed us to produce a structure-guided alignment of MRs and GPCRs and to suggest the emergence of GPCRs from Na^+^ translocating MRs [7]. Here we have updated this alignment by including the natural (not chimeric) channelrhodopsin 2, the structure of which was recently resolved [84] (ChR2, Fig. S5). As seen in Fig. S5, the Na^+^-coordinating residues of GPCRs correspond to the ion-coordinating residues from both NR and ChR2. The pivot Trp6.48 residue of the conserved WxP motif remains the only conserved residue between GPCRs and MRs, in agreement with the movement of the cytoplasmic half of helix 6 as the major conformational change both in MRs [85–87] and GPCRs [2, 9, 10, 20]. In addition to the previously reported similarities with NR, the updated alignment shows that Asp2.50 of GPCRs corresponds to a glutamate residue in helix 2 of ChR2, whereas the strictly conserved Trp4.50 of GPCRs is matched by a Trp residue in ChR2 (Fig. S5). The conservation patterns for functionally important residues between NR and GPCRs, on one hand, and ChR2 and GPCRs, on the other hand, overlap only partially (Fig. S5). It appears that the Na^+^-translocating, MR-type ancestors of GPCRs could have combined features of NR and ChR2.

As discussed previously [7], the original Na^+^-translocating MR could have evolved into an ancestral GPCR after losing its retinal moiety. A bound Na^+^ ion could stabilize the helical bundle after the loss of the retinal. Formation of a permanent binding site for the Na^+^ ion within an MR is quite feasible. Balashov and colleagues, by substituting Glu for Asp251 in the NR of *Gillisia limnaea* (which approx. corresponds to the Na^+^ ligand Asn7.45 of GPCRs, see Fig. 1, 4, S4 and Table 3), were able to create a high-affinity binding site for the Na^+^ ion in the middle of the membrane [88].

The structural similarity between GPCRs and ChR2 implies possible similarities in their mechanisms. It was shown that photoactivation of ChR was coupled not only with the motion of helix 6, but also with a rotation of helix 2 [87]. Hence, the functional mobility of helix 2, which carries the LxxxD motif, may also have been inherited from MRs.

The elements of above-described coupling machinery, including an Asp residue in the middle of the helix 2, could be seen already in representative GPCR-like proteins of early-branching eukaryotes, such as Protozoa, primitive Metazoa and plants, which are aligned in Fig. 6. Here, an Asp residue in the position that corresponds to Asp2.50 of class A GPCRs can be considered an indication of the presence of a Na^+^-binding site. Otherwise, maintaining an Asp residue in the middle of the membrane would be energetically costly and this Asp would be prone to be lost in the course of evolution, as discussed below. As follows from Fig. 6, early-branching GPCR-like proteins with a counterpart of Asp2.50 often have potential Na^+^ ligands also in the positions of other Na^+^ ligands of class A GPCRs.

The suggested key role of Leu2.46 in controlling GPCR activation is supported not only by its already mentioned high degree of conservation within class A GPCRs (90%, see [36, 64] and Table 3), but also by its conservation within all GPCR-like proteins, even those lacking Asp2.50 (Fig. 6), which is unusual for a hydrophobic residue and indicates that the shape of the side chain of Leu2.46 is particularly important. Trp6.48, the only residue that is conserved between MRs and GPCRs (Fig. S5), is highly but not universally conserved within GPCRs: it is replaced by Tyr in some Alveolata and by Phe in the protease-activated receptor 1 (PAR1) of Metazoa. In addition to Leu2.46, hydrophobic/aromatic residues are well conserved in the positions 3.40, 3.46, 6.40 and 6.44, which correspond to the hydrophobic core of class A GPCRs, see Table 3 and *cf.* Fig. 1D and 6. Residues Asn7.49 and Tyr7.53 from the NPxxY motif are all highly conserved. Finally, the characteristic motif DRY of the TM3 is absent in plants and most protozoa.

Hence, sequence comparisons of animal class A GPCRs with evolutionarily oldest, Na^+^-binding, GPCR-like proteins (Fig. 6) support our suggestion on the importance of Leu2.46, Asp2.50, Trp6.48, Asn7.49, Tyr/Phe7.53 and the set of tightly packed, conserved hydrophobic residues for the function of Na^+^-binding GPCRs and, specifically, for the coupling between their activation and Na^+^ translocation. In contrast, the DRY motif is absent from the sequences of GPCRs from plants and Alveolata and appears to be a somewhat later acquisition.

Our inspection showed that eukaryotic genomes contain numerous GPCR-like sequences, both with and without counterparts of Asp2.50. The latter sequences (some are shown in Fig. 6) most likely belong to GPCRs that cannot bind the Na^+^ ion. Presence of other Na^+^ ligands in many such sequences suggests that they could have lost their Na^+^ binding capability in the course of their evolution. Loss of Na^+^ binding in the course of GPCR evolution would not necessarily lead to the loss of function; the ability to shift between the active and inactive conformation in response to agonist binding could still be retained. Those residues that appear to form the mechanistic core of the coupling/activating mechanism in GPCRs are mostly conserved also in those GPCRs lacking a counterpart of Asp 2.50. Even without a bound Na^+^ ion, a GPCR could be driven by voltage if its activation is coupled with proton translocation across the membrane [89], e.g. via former Na^+^ ligands. However, in the absence of a Na^+^-binding site, such GPCRs would be unable to (i) suppress the noise by binding a Na^+^ ion, (ii) exploit the concentration gradient of Na^+^ ions for increasing their sensitivity and (iii) boost their selectivity by specifically amplifying the signal in response to endogenous agonists.

**Figure 6.**
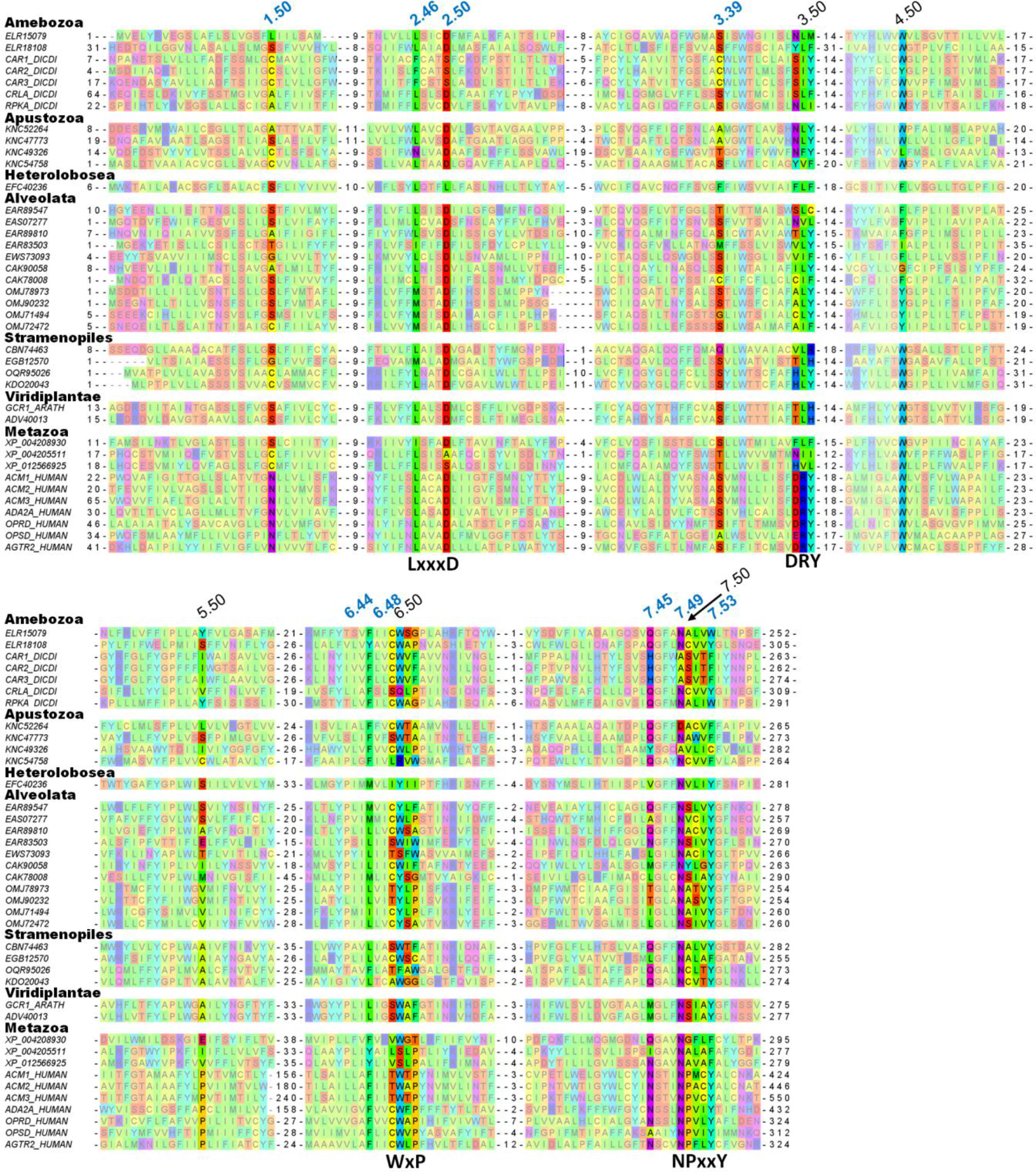
Alignment of diverse GPCRs with human class A GPCR. The top line shows Ballesteros–Weinstein numbering of the residues [17, 18] as given in GPCRdb [67] for the Class A GPCRs. Generic numbers of residues involved in the Na^+^ ion binding pocket are shown in blue. Sequences are listed under their GenBank, UniProt or RefSeq accessions and are as follows: Amoebozoa: phosphatidylinositol 4-phosphate 5-kinase protein from *Acanthamoeba castellanii* (GenBank: ELR15079); cAMP receptor protein from *Acanthamoeba castellanii* (GenBank: ELR18108); cAMP receptor 1 from *Dictyostelium discoideum* (UniProt: CAR1_DICDI); cAMP receptor 2 from *Dictyostelium discoideum* (UniProt: CAR2_DICDI); cAMP receptor 3 from *Dictyostelium discoideum* (UniProt: CAR3_DICDI); cAMP receptor-like protein from *Dictyostelium discoideum* (UniProt: CRLA_DICDI), and G-protein-coupled receptor family protein from *Dictyostelium discoideum* (UniProt: RPKA_DICDI). Apustozoa: hypothetical protein AMSG_01092 from *Thecamonas trahens* (GenBank: KNC52264); hypothetical protein AMSG_04000 from *Thecamonas trahens* (GenBank: KNC47773); hypothetical protein AMSG_02398 from *Thecamonas trahens* (GenBank: KNC56428); PPK-1 protein from *Thecamonas trahens* (GenBank: KNC49326); hypothetical protein AMSG_01609 from *Thecamonas trahens* (GenBank: KNC54758). Heterolobosea: predicted protein NAEGRDRAFT_72027 from *Naegleria gruberi* (GenBank: EFC40236). Alveolata: G protein coupled glucose receptor from *Tetrahymena thermophila* (GenBank: EAR89547); 7TM secretin family protein from *Tetrahymena thermophila* (GenBank: EAS07277); 7TM secretin family protein from *Tetrahymena thermophila* (GenBank: EAR89810); G protein coupled glucose receptor from *Tetrahymena thermophila* (GenBank: EAR83503); cAMP receptor from *Tetrahymena thermophila* (GenBank: EWS73093); unnamed protein product from *Paramecium tetraurelia* (GenBank: CAK90058); hypothetical protein from *Paramecium tetraurelia* (Genbank: CAK78008); hypothetical protein SteCoe_21100 from *Stentor coeruleus* (GenBank: OMJ78973); hypothetical protein SteCoe_30276 from *Stentor coeruleus* (GenBank: OMJ71494); hypothetical protein SteCoe_7430 from *Stentor coeruleus* (GenBank: OMJ90232); hypothetical protein SteCoe_29065 from *Stentor coeruleus* (GenBank: OMJ72472). Stramenopiles: G-protein coupled receptor from *Ectocarpus siliculosus* (GenBank: CBN74463); hypothetical protein AURANDRAFT_4432 from *Aureococcus anophagefferens*, partial (GenBank: EGB12570); hypothetical protein ACHHYP_00504 from *Achlya hypogyna* (GanBank: OQR95026); hypothetical protein SPRG_14191 from *Saprolegnia parasitica* (GenBank: KDO20043). Viridiplantae: G-protein coupled receptor 1 from *Arabidopsis thaliana* (UniProt: GCR1_ARATH); G protein coupled receptor from *Oryza sativa* (GenBank: ADV40013). Metazoa: predicted probable G-protein coupled receptor 157 from *Hydra vulgaris*, partial (NCBI RefSeq: XP_004208930); predicted cAMP receptor-like protein A from *Hydra vulgaris* (NCBI RefSeq: XP_004205511); predicted G-protein coupled receptor 1-like protein from *Hydra vulgaris* (NCBI RefSeq: XP_012566925); human muscarinic acetylcholine receptor M_1_ (UniProt: ACM1_HUMAN); human muscarinic acetylcholine receptor M_3_ (UniProt: ACM3_HUMAN); human α_2A_ adrenergic receptor (UniProt: ADA2A_HUMAN); human δ-type opioid receptor (UniProt: OPRD_HUMAN); human visual rhodopsin (UniProt: OPSD_HUMAN).

The existence of both Na^+^-dependent and Na^+^-independent GPCRs deserves comparison with rotary ATP synthases, which can be driven either by protons or Na^+^ ions [90]. While Na^+^-translocating ATP synthases are found only in some, mostly anaerobic prokaryotes [91], a comparative analysis has indicated their evolutionary primacy [79]. Apparently, in most lineages, the ability to bind and translocate Na^+^ ions got lost; these enzymes, however, retained the ability to translocate protons, which became particularly beneficial after the oxygenation of atmosphere [92]. As already mentioned, holding a Na^+^ ion in the middle of the membrane is structurally rather demanding. Therefore, it is tempting to speculate that more Na^+^-dependent GPCRs would be seen in organisms that need particularly sensitive and selective receptors for the active exploration of their environment. Indeed, Na^+^-dependent GPCRs are abundant not only in mammalian genomes, but also in genomes of primitive organisms known for their active behavior, such as free swimming, single-celled ciliates *Tetrahymena, Paramecium*, or *Stentor* (Fig. 6). For instance, the genome of *Stentor coeruleus* contains dozens of GPCR-encoding genes with a full set of Na^+^ ligands.

The Ballesteros-Weinstein nomenclature [17, 18] used in this work attributes the index "50" to the amino acid residue that is the most conserved in each helix among class A GPCRs. Comparison of Table 3 with the multiple alignment in Fig. 6 shows that the “50^th^” residues, while strictly conserved among class A GPCRs, are often not conserved within a broader set of Na^+^-binding GPCRs. Proline residues 5.50, 6.50 and 7.50, which are strictly conserved within class A GPCRs, are not conserved in receptors of primitive organisms (Fig. 6). It appears that the acquisition of additional proline residues in transmembrane helices could contribute to the success and proliferation of class A GPCRs. These proline residues could form the mechanistic scaffold of class A GPCRs, stabilize the protein fold and serve as pivots upon receptor activation. The existence of such proline scaffold would relieve the steric constrains on other residues and make class A GPCRs more prone to mutations and, hence, more evolutionarily adaptable.

In sum, Na^+^-dependent GPCRs, after their emergence in primitive eukaryotes, could be fine-tuned by successive mutations to perform diverse signaling functions. Those mutations would affect their sensitivity, signal-to-noise ratio, chemical selectivity etc. Furthermore, mutations could affect even the voltage/current profiles (where current corresponds to the signal propagation) and determine whether the particular receptor would be sensitized by membrane potential (as the majority of studied GPCRs, see Fig. 3A, 3B and Table S1) or hemmed by it (such as M_1_ receptor, see Fig. 3C and Table S1).

## Conclusions

Na^+^-binding GPCRs are splendid molecular sensors that utilize the energy of the transmembrane sodium potential to increase their (i) sensitivity; (ii) signal-to-noise ratio, and (iii) chemical selectivity. The gift of harnessing energy, in conjunction with high adaptability, might explain the presence of about 700 class A GPCR-coding genes in the human genome.

## Methods

### Model of Na^+^ translocation by class A GPCRs

Here we present the solution of the model of GPCR activation. The model implies two possible operation modes differing in the Na^+^ ion behavior upon the receptor activation: in the carrier-on mode 1, the cation barges through the membrane into the cytoplasm, while in the carrier-off mode 2 the cation returns into the extracellular space. In each mode, the system is characterized by 8 possible states of the receptor (Fig. 2) whose probabilities are defined as P_1_, …, P_8_ (Table 4).

**Table 4.** Probability coefficients for the model of GPCR activation

| Receptor state | Probability | Relative probabilities (*l*) |
| --- | --- | --- |
| Inactive receptor | P<sub>1</sub> | 1 |
| Inactive receptor with Na | P<sub>2</sub> | M·[Na]<sup>out</sup> |
| Inactive receptor with agonist | P<sub>3</sub> | N·[A] |
| Inactive receptor with agonist and Na | P<sub>4</sub> | γ·M[Na]<sup>out</sup>·N·[A] |
| Active receptor | P<sub>5</sub> | F<sub>model</sub>L<sup>a</sup> |
| Active receptor with Na | P<sub>6</sub> | α·L·M·[Na]<sup>model</sup> |
| Active receptor with agonist | P<sub>7</sub> | β·L·N·[A] |
| Active receptor with agonist and Na | P<sub>8</sub> | α·β·γ·δ·L·M·[Na]<sup>model</sup>·N·[A] |

Here, [Na]^model^ is the Na concentration available for the receptor in the active state: [Na]^in^ in the carrier-on mode and [Na]^out^ in the carrier-off mode. F_*model*_ is the electrostatic term of Na^+^ ion translocation, which is equal to *F*_2_ (see Eq. 2) in the carrier-on mode and equal to *F*_1_ (see Eq. 1) in the carrier-off mode. The forward and backward transitions between the states *m* and *n* occur at different rates *l*_1*n*_ and *l*_2*m*_; the transitions between the inactive (R) and active (R*) states of the receptor are the slowest in the system. The principle of detailed balance was applied separately to all inactive and all active receptor states, whereas the transitions between active and inactive states are treated as thermodynamically nonequilibrium. In the stationary state, the cumulative forward and backward transition probabilities between active and all inactive states match each other:

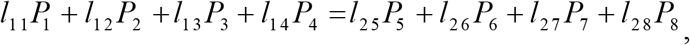

where probabilities P_1-4_ correspond to all inactive states, P_5-8_ – to all active states; *l*_1*m*_ and *l*_2*n*_ are the respective transition rate constants (see Table 1). The transition rate constants are not independent, they satisfy the thermodynamic relationship *k*_*forward*_ / *k*_*back*_ = e^−ΔG/RT^, so the latter equation can be rewritten as following:

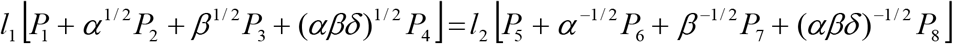

where *l*_1_ = *l*_15_ and *l*_2_ = *l*_25_. The effect of the membrane potential on cation translocation was accounted for the equilibrium constants of Na^+^ binding in both active (*F*_1_ = exp[*θ*Δ*ψF*/*RT*]) and inactive (*F*_2_ = exp[(1−*θ*)Δ*ψF* / *RT*]) states, where *θ* is the depth of the Na^+^-binding site, Δ*ψ* is the transmembrane electric potential, and *F* is the Faraday constant. This leads to the relation:

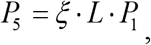

where

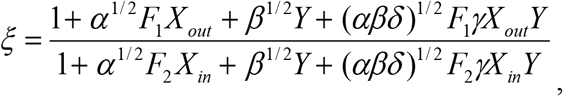

and *L* = *l*_1_/*l*_2_. In the latter equation we have used *X*_*in* / *out*_ = *M* · [*Na*^+^]_*in* / *out*_ and, *Y* = *N* · [*agonist*] respectively.

To these equations we must add the probabilities normalization requirement:

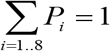

and resulting set of equations provides a solution to our model.

Suggested models for GPCR activation were implemented as Matlab R2017a functions [93]. Experimental data were obtained from respective publications and fitted with model functions using the Matlab' “lsqcurvefit” function. During the fit, the allosteric coefficients α-δ and the depth of the Na^+^-binding site were kept constant. The goal was to find coefficients M (Na^+^-binding constant) and N (agonist-binding constant) that provided the best fit of the experimental data (two sets of data points obtained at different membrane potential) with the two model curves calculated with identical parameters except for the membrane potential values.

### Structure analysis

Structure superposition and visualization were performed with PyMol v 1.7 [72] and YASARA [94]. Structure analyses were performed with the WHAT IF [95] subset of the YASARA Twinset. Cavities and caves were calculated using the method by Voorintholt et al [96] with a spherical probe of the 1.4 Å radius. These were visualized using a 1 Å resolution grid. Structures were superposed using the method of Vriend and Sander [97]. Rotamer distributions were calculated using the method of Chinea et al. [70]. Briefly, the rotamer distribution software searches the PDB for stretches of five residues that (i) have a very similar backbone as observed in the local structure and (ii) have the same middle residue as the local structure. The obtained database stretches is then superposed on the local structure, but only the side chain of the middle residue is shown.

## Abbreviations

cAMP: 3'5'-cyclic adenosine monophosphate
GPCR: G-protein coupled receptor
MR: microbial rhodopsin
RMSD: root-mean-square deviation
TM: transmembrane
NR: Na-translocating microbial rhodopsin
ChR2: channelrhodopsin 2

## Acknowledgements

The authors would like to thank Profs. V. Katritch, H.-J. Steinhoff and H. Vogel for encouragement and very useful discussions. Helpful advises of Dr. N.P. Isaev are greatly appreciated. The calculations were done using the equipment of the shared research facilities of HPC computing resources at Lomonosov Moscow State University supported by its Development Program. This study was supported by the *Deutsche Forschungsgemeinschaft*, Federal Ministry of Education and Research of Germany, the EvoCell Program of the Osnabrueck University (AYM), the *Ostpartnerschaftenprogramm* of the German Academic Exchange Service (*DAAD*), the Russian Government contract (AAAA-A19-119012890064-7), and the 14-50-00029 grant from the Russian Science Foundation (DNS). MYG is supported by the Intramural Research Program of the NIH at the National Library of Medicine.

## Author contributions

AYM designed the study. AYM, DAC and DNS developed the model. DNS calculated the model and fitted experimental data. DNS, AYM, MYG and GV performed the structural analysis. DNS, MYG and AYM performed the evolutionary analysis. DNS, DAC, MYG, GV and AYM wrote the paper.

## Competing interests

Authors confirm that there have been no involvements that might raise the question of bias in the work reported or in the conclusions, implications, or opinions stated.

