## Supplementary Table S1 and Supplementary Figures S1-S5 for "Class A GPCRs use the membrane potential to increase their sensitivity and selectivity"

**Supplementary Table S1. Experimental data on voltage-sensitive activation of GPCRs**.

| Receptor | Agonist | Agonist characterization | Voltage effect on activation | Method of activity measurement | Reference |
|---|---|---|---|---|---|
| M2 muscarinic receptor | acetylcholine | full agonist (endogenous) | enhanced | GIRK currents | [1, 2] |
| M2 muscarinic receptor | oxotremorine | full/strong agonist | enhanced | | |
| M1 muscarinic receptor | acetylcholine | full agonist (endogenous) | decreased | | |
| M2 muscarinic receptor | acetylcholine | full agonist (endogenous) | enhanced | GIRK currents | [3] |
| M2 muscarinic receptor | acetylcholine | full agonist (endogenous) | enhanced | ACh-activated $K^+$ current | [2, 4] |
| M2 muscarinic receptor | pilocarpine | partial agonist | decreased | | |
| M2 muscarinic receptor | acetylcholine | full agonist (endogenous) | enhanced | ACh-activated $K^+$ current | [2, 5] [6] |
| M2 muscarinic receptor | pilocarpine | partial agonist | decreased | | |
| M2 muscarinic receptor | bethanechol | Agonist (low affinity) | no effect | | |
| M1 muscarinic receptor | carbachol | full/strong agonist | decreased | FRET–based assays | [2, 7] |
| M3 muscarinic receptor | carbachol | full/strong agonist | enhanced | | |
| mGluR3 glutamate receptor | glutamate | full agonist (endogenous) | enhanced | $K^+$ currents and $Cl^-$ currents | [8] |
| mGluR1a glutamate receptor | glutamate | full agonist (endogenous) | decreased | | |
| $\alpha_{2A}$-AR adrenergic receptor | noradrenaline | full agonist (endogenous) | enhanced | FRET–based assays | [9] |

| | | | | | |
|---|---|---|---|---|---|
| $\beta_1$-AR adrenergic receptor | isoprenaline | full agonist | enhanced | FRET–based assays | [10, 11] |
| $\beta_1$-AR adrenergic receptor | adrenaline | full agonist (endogenous) | enhanced | | |
| dopamine D2L receptor | dopamine | full agonist (endogenous) | enhanced | GIRK currents | [12, 13] |
| dopamine D2L receptor | quinpirole | full agonist | enhanced | | |
| dopamine D2S receptor | dopamine | full agonist (endogenous) | | GIRK currents | [14] |
| dopamine D2S receptor | dopamine | full agonist (endogenous) | enhanced | GIRK currents | [12, 15] |
| dopamine D2S receptor | *p*-tyramine | partial agonist | decreased | | |
| dopamine D2S receptor | *m*-tyramine | partial agonist | decreased | | |
| dopamine D2S receptor | phenylamine | partial agonist | decreased | | |
| dopamine D2S receptor | S-5-OH-DPAT | full agonist | no effect | | |
| dopamine D2S receptor | R-5-OH-DPAT | full/strong agonist[12] | no effect | | |
| dopamine D2S receptor | R-7-OH-DPAT | full/strong agonist | no effect | | |
| dopamine D2S receptor | RIS-OH-DPAT | full/strong agonist[12] | no effect | | |

The effectiveness of a signaling molecule in the receptor activation does not necessarily reflect the affinity of that molecule to the receptor. Some partial agonists have a high binding affinity but induce lower receptor activity even when added in saturating amounts [2, 15]. Such molecules are believed to stabilize the receptor in an intermediate state and/or stimulate the transition to the active state with lower probability than full agonists.

**Supplementary Figures**

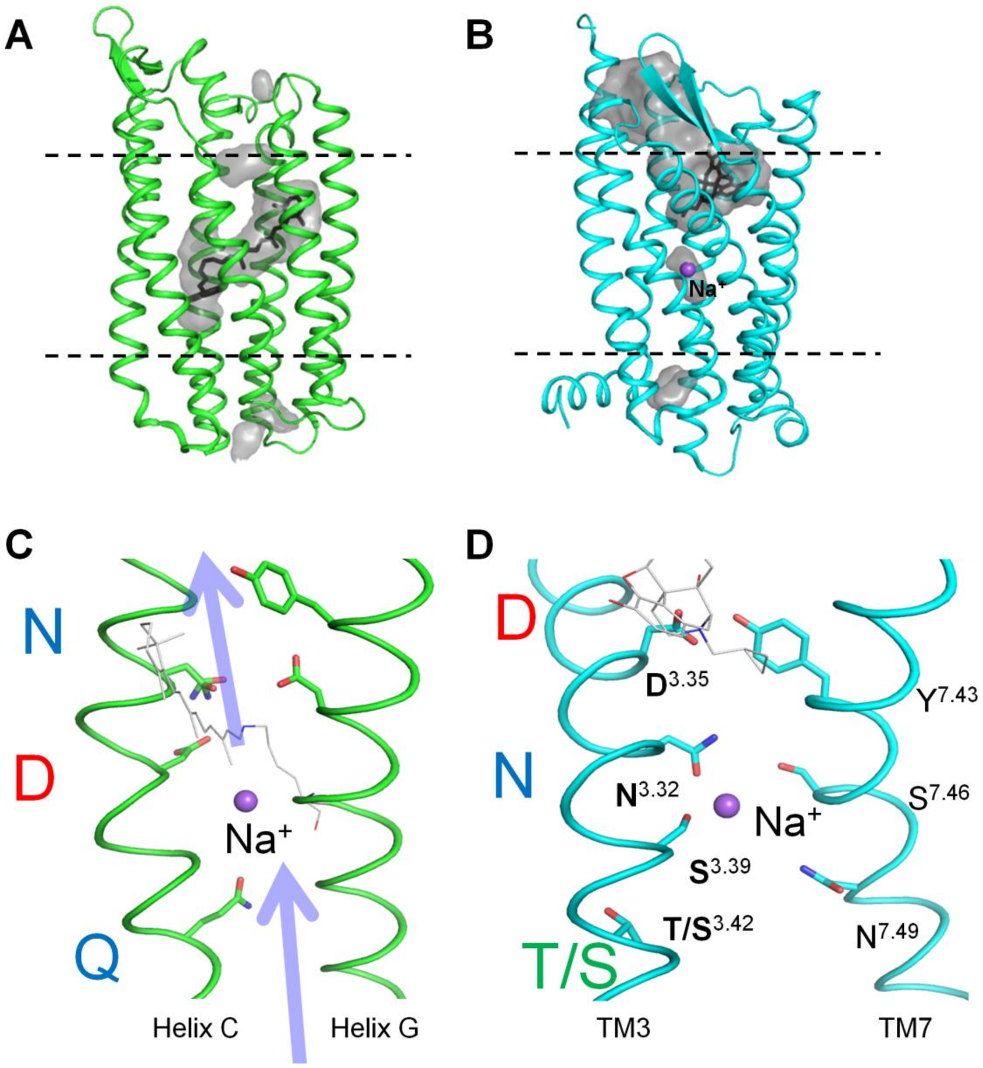


**Figure S1. Structural and functional similarities between microbial rhodopsins (MRs) and GPCRs.** A. Structure of sodium-translocating rhodopsin KR2 (PDB 4XTL); the protein is shown as a cartoon, Lys255 and retinal are shown as black sticks, cavities inside the protein are shown as gray volumes. B. Structure of the human δ-opioid receptor δ-OR (PDB 4N6H); the protein is shown as a cartoon, antagonist (naltrindole) is shown as black sticks, cavities inside the protein are shown as gray volumes. Cavities are depicted as defined by CASTp service http://sts.bioe.uic.edu/castp/ [16]. C. Signature residues involved in cation translocation by $Na^+$-transporting rhodopsin KR2 (panel A). Signature residues of the NDQ motif are shown as sticks, Lys255 and retinal are shown in gray. D. Corresponding residues in helices TM3 and TM7 in δ-OR (panel B), antagonist (naltrindole) shown in gray, residue numbering is according to Ballesteros-Weinstein [17].

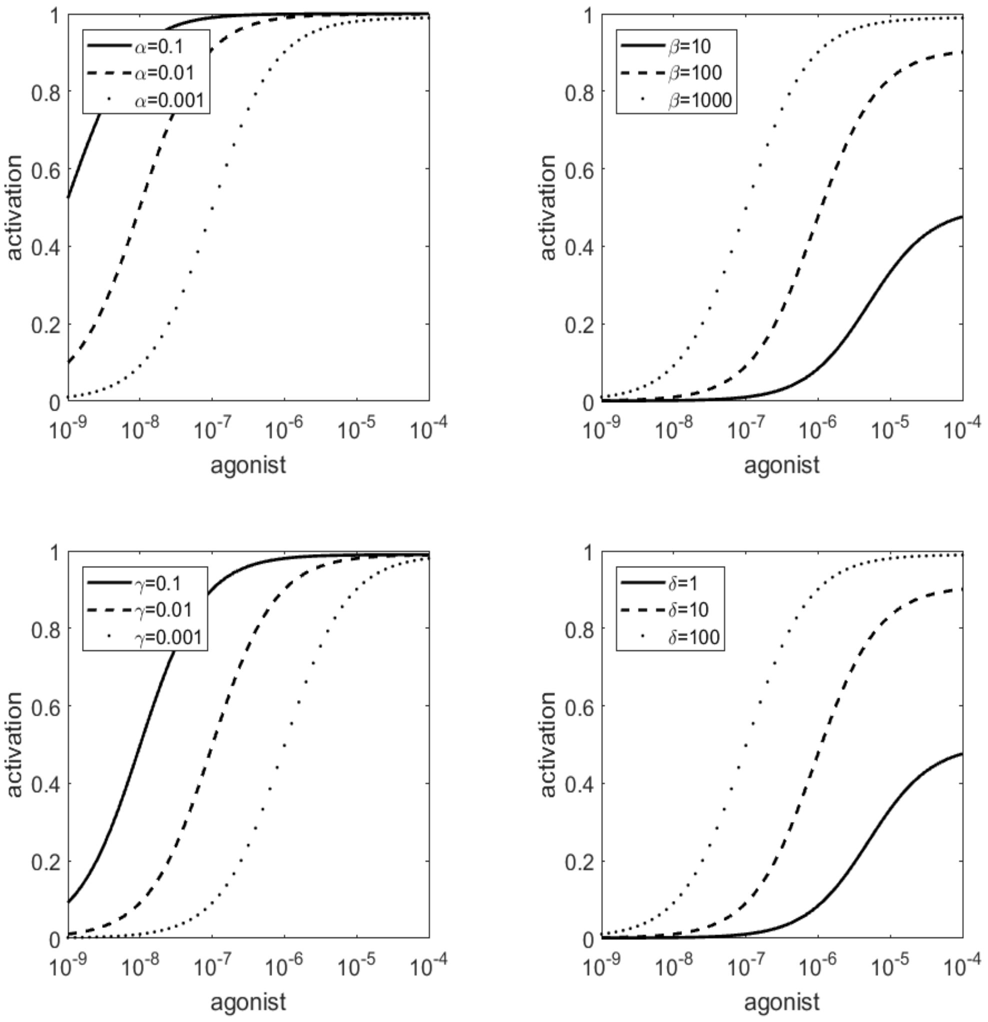


**Figure S2. Effects of allosteric parameters α,β,γ and δ on the activation curves.** Dependence of the GPCR activation on the coefficient α, intrinsic efficacy of sodium (A); coefficient β, intrinsic efficacy of the agonist (B); coefficient γ, binding cooperativity between sodium and the agonist (C), and coefficient δ, activation cooperativity between sodium and the agonist (D). In each panel, one of parameters was varied while the values of other parameters were taken from Table 1. The impact of the membrane voltage was not taken into account in the calculations.

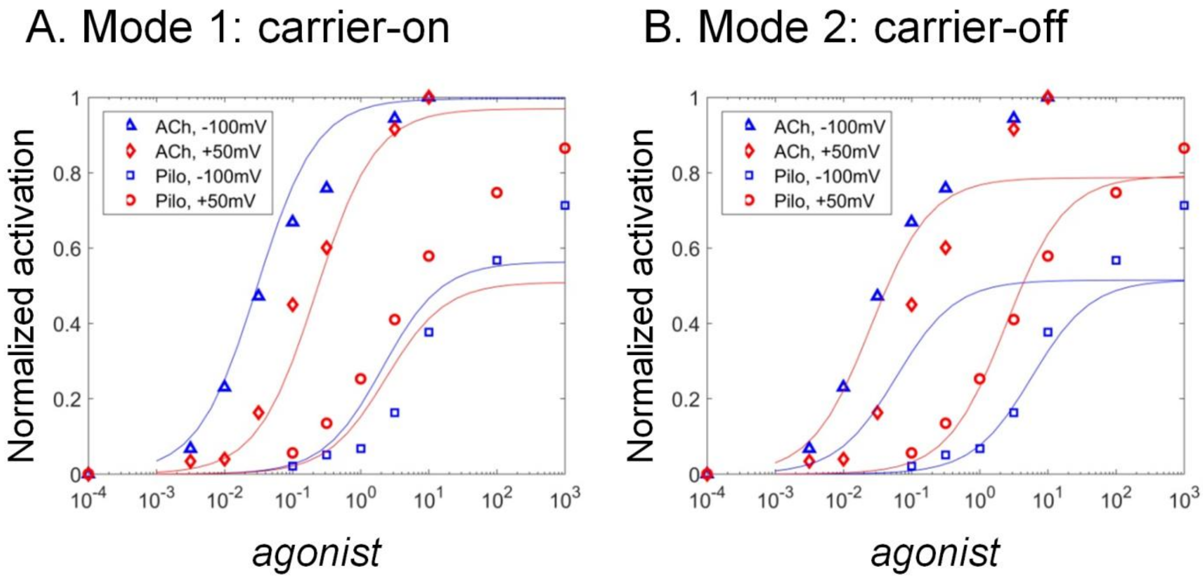


**Figure S3. Fitting experimental data of voltage-sensitive muscarinic acetylcholine receptor $M_2$ activation with the suggested kinetic models**. Concentration-response curves, as obtained for the full endogenous agonist acetylcholine (ACh) and partial synthetic agonist pilocarpine (Pilo), measured at different magnitudes of membrane potential were fitted with model 1 (A) and model 2 (B). Experimental data from [4].

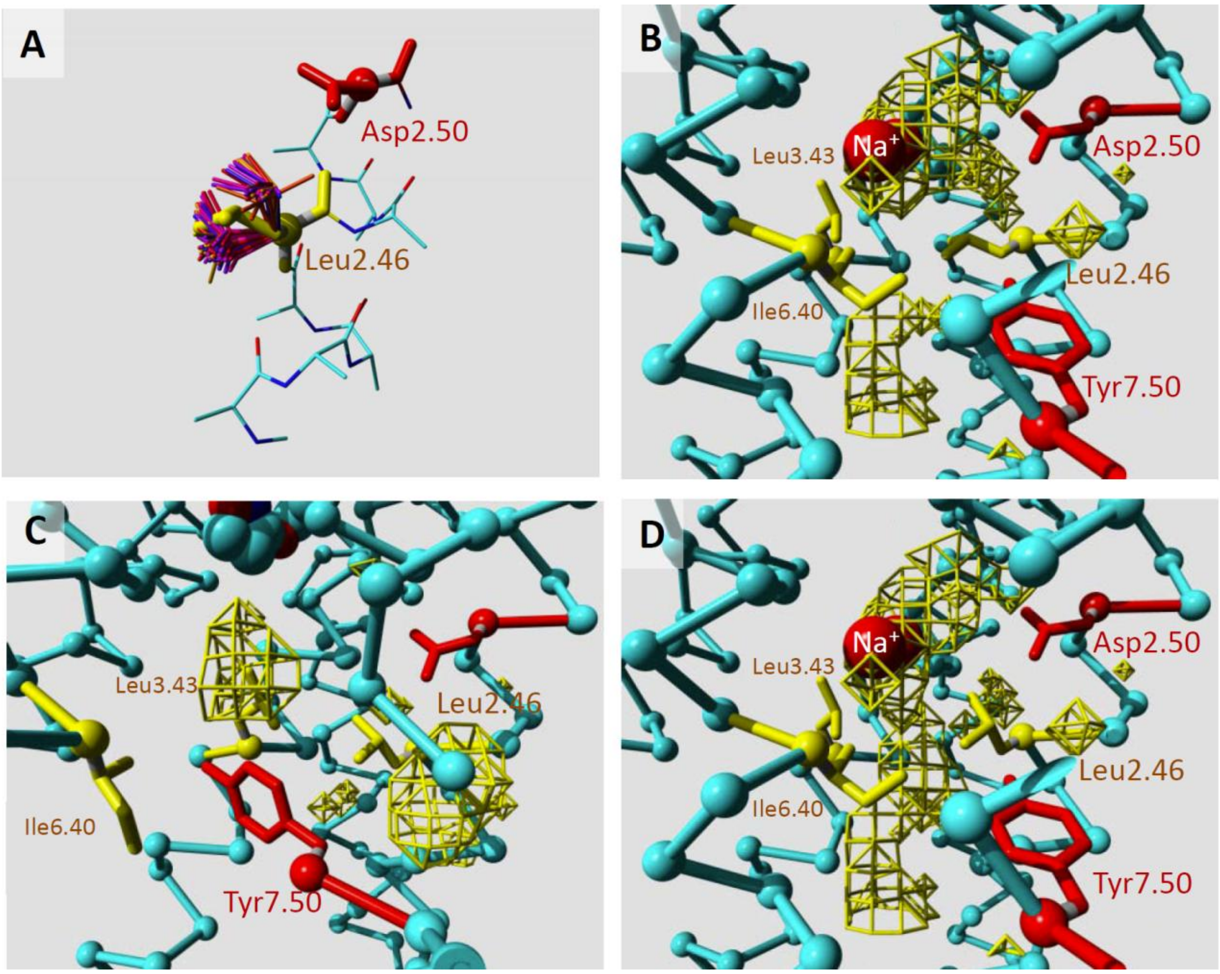


**FigureS4. Rotamers of Leu2.46 in the inactive and inactive conformations of muscarinic acetylcholine receptor $M_2$.**

A, Analysis of Leu2.46 rotamers, shown on the example of PDB 3UON. The section of Helix 2 is shown coloured by atom type. The Asp2.50 side chain is shown in red. Leu2.46 as in the 3UON structure is shown in yellow. The preferable rotamers for leucine at position 2.46 are shown in different shades of orange, blue, and red. B-D, The $Na^+$ binding site in the structure of the inactive receptor, PDB 3UON (B); the active receptor (PDB 4MQT) (C), and the inactive receptor (PDB 3UON) with Leu2.46 moved into upper rotamer. C-alpha traces are shown in light-blue (some residues in the front that obscured the view have been removed from the plots). Side chains of Asp2.50 and Tyr7.50 are shown as red stick models. Cavities large enough to hold water molecules are represented by yellow mesh. A piece of the ligand in the inactive structure (B and D, PDB 3UON) is seen in purple. The side chains of the three aliphatic residues Leu2.46, Leu3.43, and Ile6.40, are shown as yellow stick models.

**Fig. S5. Multiple sequence alignment of transmembrane segments in GPCRs and MRs.**

Sequences used in the alignment (listed under their UniProt IDs):

**GPCRs:**

| | |
|---|---|
| ACM1_HUMAN | Human muscarinic acetylcholine receptor M1 |
| ACM3_RAT | Rat muscarinic acetylcholine receptor M3 |
| ADRB1_MELGA | Turkey β-1 adrenergic receptor |
| DRD2_HUMAN | Human $D_2$ dopamine receptor |
| HRH1_HUMAN | Human histamine H1 receptor |
| OPRD_HUMAN | Human δ-type opioid receptor |
| OPRM_MOUSE | Mouse μ-type opioid receptor |
| OPRX_HUMAN | Human nociceptin receptor |
| CCR2_HUMAN | Human C-C chemokine receptor type 2 |
| CCR5_HUMAN | Human C-C chemokine receptor type 5 |
| AA1R_HUMAN | Human adenosine receptor A1 |
| AA2AR_HUMAN | Human adenosine receptor A2a |
| OPSD_HUMAN | Human visual rhodopsin |
| OPSD_TODPA | Squid visual rhodopsin |
| PAR1_HUMAN | Human proteinase-activated receptor 1 |
| PAR2_HUMAN | Human proteinase-activated receptor 2 |

**MRs**

| | |
|---|---|
| Q8RUT8_CHLRE | Channelrhodopsin 2 from Chlamydomonas reinhardtii |
| W8VZW3_9FLAO | Chloride pumping rhodopsin from Nonlabens marinus |
| N0DKS8_9FLAO | Sodium pumping rhodopsin from Dokdonia eikasta |
| BACH_HALS3 | Halorhodopsin from Halobacterium salinarum |
| Q2S2F8_SALRD | Xanthorhodopsin from Salinibacter ruber |
| S5DM51_9ACTN | Bacteriorhodopsin from Candidatus Actinomarina minuta |
| BACS2_NATPH | Sensory rhodopsin-2 from Natronomonas pharaonis |
| BACR1_HALC1 | Archaerhodopsin-1 from Halorubrum chaoviator |
| BACR1_HALMA | Bacteriorhodopsin-I from Haloarcula marismortui |
| BACR1_HALWC | Bacteriorhodopsin-I from Haloquadratum walsbyi |
| BACR_HALSA | Bacteriorhodopsin from Halobacterium salinarum |
| BACR2_HALS2 | Archaerhodopsin-2 from Halobacterium sp. |
| BACS1_HALSA | Sensory rhodopsin-1 from Halobacterium salinarum |
| Q93WP2_CHLRE | Archaeal-type opsin 1 from Chlamydomonas reinhardtii |
| I4DST7_9EURY | Deltarhodopsin from Haloterrigena thermotolerans |

Alignment of GPCRs was taken from the GPCRdb web service [18], alignment of MRs was constructed with T-Coffee[19], two alignments were merged using the structure superposition of human δ-opioid receptor (PDB 4N6H) and channelrhodopsin 2 from *Chlamydomonas reinhardtii* (PDB 6EIG), as described in [20].

| **Helix 1/A** | **1.50** |
|---|---|
| ---GPCRs--- | |
| ACM1_HUMAN/22-53 | PWQVAFIGITTGLLSLATVTG**N**LLVLISFKVN |
| ACM3_RAT/64-95 | IWQVVFIAFLTGFLALVTIIG**N**ILVIVAFKVN |
| ADRB1_MELGA/38-69 | QQWEAGMSLLMALVVLLIVAG**N**VLVIAAIGRT |
| DRD2_HUMAN/31-62 | RPHYNYYATLLTLLIAVIVFG**N**VLVCMAVSRE |
| RHR1_HUMAN/29-55 | -----PLVVVLSTICLVTVGL**N**LLVLYAVRSE |
| OPRD_HUMAN/46-77 | LALAIAITALYSAVCAVGLLG**N**VLVMFGIVRY |
| OPRM_MOUSE/65-96 | MVTAITIMALYSIVCVVGLFG**N**FLVMYVIVRY |
| OPRX_HUMAN/48-79 | LGLKVTIVGLYLAVCVGGLLG**N**CLVMYVILRH |
| CCR2_HUMAN/39-70 | QIGAQLLPPLYSLVFIFGFVG**N**MLVVLILINC |
| CCR5_HUMAN/27-58 | QIAARLLPPLYSLVFIFGFVG**N**MLVILILINC |
| AA1R_HUMAN/6-37 | SAFQAAYIGIEVLIALVSVPG**N**VLVIWAVKVN |
| AA2AR_HUMAN/3-34 | IMGSSVYITVELAIAVLAILG**N**VLVCWAVWLN |
| OPSD_HUMAN/34-65 | PWQFSMLAAYMFLLIVLGFPI**N**FLTLYVTVQH |
| OPSD_TODPA/31-62 | DAVYYSLGIFIGICGIIGCGG**N**GIVIYLFTKT |
| PAR1_HUMAN/99-130 | SWLTLFVPSVYTGVFVVSLPL**N**IMAIVVFILK |
| PAR2_HUMAN/72-103 | KLTTVFLPIVYTIVFVVGLPS**N**GMALWVFLFR |
| ----MRs--- |  |
| Q8RUT8_CHLRE/50-80 | TASNVLQWLAA-GFSILLLMF**Y**AYQTWKSTCG |
| W8VZW3_9FLAO/16-46 | EFIDHLLTMGV-GVHFAALIF**F**LVVSQFVAPK |
| N0DKS8_9FLAO/26-56 | QFTSHILTLGY-AVMLAGLLY**F**ILTIKNVDKK |
| BACH_HALS3/27-57 | ALLSSSLWVNV-ALAGIAILV**F**VYMGRTIRPG |
| Q2S2F8_SALRD/14-44 | SLVFNMFSFTV-ATMTASFVF**F**VLARNNVAPK |
| S5DM51_9ACTN/3-33 | ELTYRLFMVAT-VGMLAGTVF**L**LASSREVKPE |
| BACS2_NATPH/2-32 | VGLTTLFWLGA-IGMLVGTLA**F**AWAGRDAGSG |
| BACR1_HALC1/19-49 | RPETLWLGIGT-LLMLIGTFY**F**IVKGWGVTDK |
| BACR1_HALMA/5-35 | GSEGIWLWLGT-AGMFLGMLY**F**IARGWGETDG |
| BACR1_HALWC/14-44 | EGEGIWLALGT-IGMLLGMLY**F**IADGLDVQDP |
| BACR_HALSA/20-50 | RPEWIWLALGT-ALMGLGTLY**F**LVKGMGVSDP |
| BACR2_HALS2/17-47 | RPETLWLGIGT-LLMLIGTFY**F**IARGWGVTDK |
| BACS1_HALSA/2-32 | DAVA-TAYLGGAVALIVGVAF**V**WLLYRSLDGS |
| Q93WP2_CHLRE/89-119 | LAANILQWITF-ALSALCLMF**Y**GYQTWKSTCG |
| I4DST7_9EURY/6-36 | GPESIWLWIGT-IGMTLGTLY**F**VGRGRGVRDR |

| **Helix 2/B** | **2.46 2.50** |
| --- | --- |
| ---GPCRs--- | |
| ACM1_HUMAN/58-88 | TVNNYFLLS**L**ACA**D**LIIGTFSMN-LYTTYLLM |
| ACM3_RAT/100-130 | TVNNYFLLS**L**ACA**D**LIIGVISMN-LFTTYIIM |
| ADRB1_MELGA/74-104 | TLTNLFITS**L**ACA**D**LVMGLLVVP-FGATLVVR |
| DRD2_HUMAN/67-97 | TTTNYLIVS**L**AVA**D**LLVATLVMP-WVVYLEVV |
| HRH1_HUMAN/60-90 | TVGNLYIVS**L**SVA**D**LIVGAVVMP-MNILYLLM |
| OPRD_HUMAN/82-111 | TATNIYIFN**L**ALA**D**ALATS-TLP-FQSAKYLM |
| OPRM_MOUSE/101-130 | TATNIYIFN**L**ALA**D**ALATS-TLP-FQSVNYLM |
| OPRX_HUMAN/84-113 | TATNIYIFN**L**ALA**D**TLVLL-TLP-FQGTDILL |
| CCR2_HUMAN/75-104 | CLTDIYLLN**L**AIS**D**LLFLI-TLP-LWAHSAAN |
| CCR5_HUMAN/63-92 | SMTDIYLLN**L**AIS**D**LFFLL-TVP-FWAHYAAA |
| AA1R_HUMAN/42-72 | DATFCFIVS**L**AVA**D**VAVGALVIP-LAILINIG |
| AA2AR_HUMAN/39-69 | NVTNYFVVS**L**AAA**D**IAVGVLAIP-FAITISTG |
| OPSD_HUMAN/70-100 | TPLNYILLN**L**AVA**D**LFMVLGGFT-STLYTSLH |
| OPSD_TODPA/67-97 | TPANMFIIN**L**AFS**D**FTFSL-VNGFPLMTISCF |
| PAR1_HUMAN/135-164 | KPAVVYMLH**L**ATA**D**VLFVS-VLP-FKISYYFS |
| PAR2_HUMAN/108-137 | HPAVIYMAN**L**ALA**D**LLSVI-WFP-LKIAYHIH |
| ----MRs--- |  |
| Q8RUT8_CHLRE/89-111 | IEMVKVILE**F**F-F**E**FKNPSMLYLA-------- |
| W8VZW3_9FLAO/54-84 | SCIVMVSAG**L**ILN**S**QAVMWTDAYAYVDGSYQ- |
| N0DKS8_9FLAO/64-95 | SAVVMVSAF**L**LLY**A**QAQNWTSSFTFNEEVGRY |
| BACH_HALS3/66-97 | TLMIPLVSI**S**SYL**G**LLSGLTVGMIEMPAGHAL |
| Q2S2F8_SALRD/52-81 | SALVVFIAG**Y**HYF**R**ITSSWEAAYALQNGM--Y |
| S5DM51_9ACTN/41-62 | SALVCGIAW**Y**HYQ**K**MGASWESG---------- |
| BACS2_NATPH/40-64 | LVGISGIAA**V**AYV**V**MALGVGWVPVA------- |
| BACR1_HALC1/58-85 | TILVPGIAS**A**AYL**S**MFFGIGLTEVQVGS---- |
| BACR1_HALMA/44-71 | TILITAIAF**V**NYL**A**MALGFGLTFIEFGG---- |
| BACR1_HALWC/53-81 | TILIPAIAA**A**SYL**S**MFFGFGLTEVSLANG--- |
| BACR_HALSA/59-86 | TTLVPAIAF**T**MYL**S**MLLGYGLTMVPFGG---- |
| BACR2_HALS2/56-84 | TILVPGIAS**A**AYL**A**MFFGIGVTEVELASG--- |
| BACS1_HALSA/41-65 | LAIIPVFAG**L**SYV**G**MAYDIGTVIVN------- |
| Q93WP2_CHLRE/127-150 | TIEMIKFII**E**-YF**H**EFDEPAVIYSS------- |
| I4DST7_9EURY/45-72 | TTFITTIAA**A**MYF**A**MATGFGVTEVVVGD---- |

| **Helix 3/C** | **3.32 3.39 3.50** |
|---|---|
| ---GPCRs--- | |
| ACM1_HUMAN/95-125 | TLACDLWLAL**D**YVASNA**S**VMNLLLISFDRYF |
| ACM3_RAT/137-167 | NLACDLWLSI**D**YVASNA**S**VMNLLVISFDRYF |
| ADRB1_MELGA/111-141 | SFLCECWTSL**D**VLCVTA**S**IETLCVIAIDRYL |
| DRD2_HUMAN/104-134 | RIHCDIFVTL**D**VMMCTA**S**ILNLCAISIDRYT |
| HRH1_HUMAN/97-127 | RPLCLFWLSM**D**YVASTA**S**IFSVFILCIDRYR |
| OPRD_HUMAN/118-148 | ELLCKAVLSI**D**YYNMFT**S**IFTLTMMSVDRYI |
| OPRM_MOUSE/137-167 | NILCKIVISI**D**YYNMFT**S**IFTLCTMSVDRYI |
| OPRX_HUMAN/120-150 | NALCKTVIAI**D**YYNMFT**S**TFTLTAMSVDRYV |
| CCR2_HUMAN/110-140 | NAMCKLFTGL**Y**HIGYFG**G**IFFIILLTIDRYL |
| CCR5_HUMAN/98-128 | NTMCQLLTGL**Y**FIGFFS**G**IFFIILLTIDRYL |
| AA1R_HUMAN/77-107 | FHTCLMVACP**V**LILTQS**S**ILALLAIAVDRYL |
| AA2AR_HUMAN/74-104 | CHGCLFIACF**V**LVLTQS**S**IFSLLAIAIDRYI |
| OPSD_HUMAN/107-137 | PTGCNLEGFF**A**TLGGEI**A**LWSLVVLAIERYV |
| OPSD_TODPA/105-135 | FAACKVYGFI**G**GIFGFM**S**IMTMAMISIDRYN |
| PAR1_HUMAN/172-202 | SELCRFVTAA**F**YCNMYA**S**ILLMTVISIDRFL |
| PAR2_HUMAN/145-175 | EALCNVLIGF**F**YGNMYC**S**ILFMTCLSVQRYW |
| ----MRs--- |  |
| Q8RUT8_CHLRE/113-143 | GHRVQWLRYA**E**WLLTCP**V**ILIHLSNLTGLSN |
| W8VZW3_9FLAO/88-118 | LTFSNGYRYV**N**WMATIP**C**LLLQLLIVLNLKG |
| N0DKS8_9FLAO/102-132 | DLFNNGYRYL**N**WLIDVP**M**LLFQILFVVSLTT |
| BACH_HALS3/102-131 | -VRSQWGRYL**T**WALSTP**M**ILLALGLLADVDL |
| Q2S2F8_SALRD/86-116 | ELFNDAYRYV**D**WLLTVP**L**LTVELVLVMGLPK |
| S5DM51_9ACTN/63-92 | -SYDTGLRYV**D**WVLTVP**L**MFVEVLAVTRKGA |
| BACS2_NATPH/65-95 | ERTVFAPRYI**D**WILTTP**L**IVYFLGLLAGLDS |
| BACR1_HALC1/87-117 | MLDIYYARYA**D**WLFTTP**L**LLLDLALLAKVDR |
| BACR1_HALMA/73-103 | QHPIYWARYT**D**WLFTTP**L**LLYDLGLLAGADR |
| BACR1_HALWC/83-113 | VVDVYWARYA**D**WLFTTP**L**LLLDIGLLAGASQ |
| BACR_HALSA/88-118 | QNPIYWARYA**D**WLFTTP**L**LLLDLALLVDADQ |
| BACR2_HALS2/86-116 | VLDIYYARYA**D**WLFTTP**L**LLLDLALLAKVDR |
| BACS1_HALSA/66-96 | GNQIVGLRYI**D**WLVTTP**I**LVGYVGYAAGASR |
| Q93WP2_CHLRE/152-182 | GNKTVWLRYA**E**WLLTCP**V**ILIHLSNLTGLAN |
| I4DST7_9EURY/74-104 | ALTIYWARYA**D**WLFTTP**L**LLLDLGLLAGANR |

```
Helix 4/D                         4.50
---GPCRs------------------------------------------
ACM1_HUMAN/138-159    TPRRAALMIGLAWLVSFVLWAP
ACM3_RAT/180-201      TTKRAGVMIGLAWVISFVLWAP
ADRB1_MELGA/154-175   TRARAKVIICTVWAISALVSFL
DRD2_HUMAN/148-168    SKRRVTVMISIVWVLSFTIS-C
HRH1_HUMAN/140-160    TKTRASATILGAWFLSFL-WVI
OPRD_HUMAN/161-182    TPAKAKLINICIWVLASGVGVP
OPRM_MOUSE/180-201    TPRNAKIVNVCNWILSSAIGLP
OPRX_HUMAN/163-184    TSSKAQAVNVAIWALASVVGVP
CCR2_HUMAN/153-173    TVTFGVVTSVITWLVAVFAS-V
CCR5_HUMAN/141-161    TVTFGVVTSVITWVVAVFAS-L
AA1R_HUMAN/120-141    TPRRAAVAIAGCWILSFVVGLT
AA2AR_HUMAN/117-138   TGTRAKGIIAICWVLSFAIGLT
OPSD_HUMAN/149-170    GENHAIMGVAFTWVMALACAAP
OPSD_TODPA/148-169    SHRRAFIMIIFVWLWSVLWAIG
PAR1_HUMAN/215-235    TLGRASFTCLAIWALAIAGV-V
PAR2_HUMAN/187-207    KANIAIGISLAIWLLILLVTI-
----MRs-------------------------------------------
Q8RUT8_CHLRE/152-171 --LLVSDIGTIVWGATSAMATG
W8VZW3_9FLAO/128-148 -LILAAWGMIITGYVGQLYEVD
N0DKS8_9FLAO/142-161 --FWFSGAMMIITGYIGQFYEV
BACH_HALS3/141-157    --DIGMCVTGLAAAMTTSA---
Q2S2F8_SALRD/127-146 GFLAALMIVLGYPGEVSENA--
S5DM51_9ACTN/101-120 --WGIAATVMIGAGYYGETSAA
BACS2_NATPH/105-117   --NTVVMLAGFAGAM-------
BACR1_HALC1/127-143   --DALMIVTGLVGALSHTP---
BACR1_HALMA/113-132   --DVLMIGTGVVATLSAGSGVL
BACR1_HALWC/123-139   --DAFMIVTGLVATLTKVV---
BACR_HALSA/128-144    --DGIMIGTGLVGALTKVY---
BACR2_HALS2/126-142   --DALMIVTGLIGALSKTP---
BACS1_HALSA/106-121   --DALMIAVGAGAVVTDG----
Q93WP2_CHLRE/194-211 ----SDIGTIVWGTTAALSKGY
I4DST7_9EURY/114-130 --DVFMIGTGMIAAFAATP---
```

```
Helix 5/E                           5.50
---GPCRs----------------------------------------
ACM1_HUMAN/185-211   QPIITF-GTAMAAFYLPVTVMCTLYWRI
ACM3_RAT/227-253     EPTITF-GTAIAAFYMPVTIMTILYWRI
ADRB1_MELGA/204-230  NRAYAI-ASSIISFYIPLLIMIFVYLRV
DRD2_HUMAN/186-212   NPAFVV-YSSIVSFYVPFIVTLLVYIKI
HRH1_HUMAN/187-213   VTWFKV-MTAIINFYLPTLLMLWFYAKI
OPRD_HUMAN/210-236   DTVTKI-CVFLFAFVVPILIITVCYGLM
OPRM_HUMAN/229-255   ENLLKI-CVFIFAFIMPVLIITVCYGLM
OPRX_HUMAN/212-238   GPVFAI-CIFLFSFIVPVLVISVCYSLM
CCR2_HUMAN/199-225   NNFHTI-MRNILGLVLPLLIMVICYSGI
CCR5_HUMAN/191-217   KNFQTL-KIVILGLVLPLLVMVICYSGI
AA1R_HUMAN/176-203   SMEYMVYFNFFVWVLPPLLLMVLIYLEV
AA2AR_HUMAN/173-200  PMNYMVYFNFFACVLVPLLLMLGVYLRI
OPSD_HUMAN/200-226   NESFVI-YMFVVHFTIPMIIIFFCYGQL
OPSD_TODPA/197-223   TRSNIL-CMFILGFFGPILIIFFCYFNI
PAR1_HUMAN/268-293   AYYFSA-FSAVF-FFVPLIISTVCYVSI
PAR2_HUMAN/239-265   MFNYFL-SLAIGVFLFPAFLTASAYVLM
----MRs-----------------------------------------
Q8RUT8_CHLRE/173-197 ---VKVIFFCLGLCYGANTFFHAAKAYI
W8VZW3_9FLAO/150-176 I-AQLMIWGAVSTAFFVVMNWIVGTKIF
N0DKS8_9FLAO/164-190 L-TAFLVWGAISSAFFFHILWVMKKVIN
BACH_HALS3/161-187   RWAFYAISCAFFVVVLSALVTDWAASA-
Q2S2F8_SALRD/148-175 LFGTRGLWGFLSTIPFVWILYILFTQLG
S5DM51_9ACTN/121-145 --GSNEYWTGFVIAM-ATYVWLMRNLQA
BACS2_NATPH/123-149  RYALFGMGAVAFLGLVYYLVGPMTESA-
BACR1_HALC1/146-172  RYTWWLFSTICMIVVLYFLATSLRAAA-
BACR1_HALMA/138-164  RLVWWGISTAFLLVLLYFLFSSLSGRV-
BACR1_HALWC/142-168  RYAFWTISTISMVFLLYYLVAVFGEAV-
BACR_HALSA/147-173   RFVWWAISTAAMLYILYVLFFGFTSKA-
BACR2_HALS2/145-171  RYTWWLFSTIAFLFVLYYLLTSLRSAA-
BACS1_HALSA/124-149  KWALFGVSSIFHLSLFAYLYVIFPRV--
Q93WP2_CHLRE/212-239 VRVIFFLMGLCYGIYTFFNAAKVYIEAY
I4DST7_9EURY/133-159 RIAWWGISTGALLALLYVLVGTLSKDA-
```

| **Helix 6/F** | **6.48** |
|---|---|
| ---GPCRs--- | ----------------------------- |
| ACM1_HUMAN/363-387 | -AARTLSAILLAFILT**W**TPYNIMVLV |
| ACM3_RAT/488-512 | -AAQTLSAILLAFIIT**W**TPYNIMVLV |
| ADRB1_MELGA/288-312 | -ALKTLGIIMGVFTLC**W**LPFFLVNIV |
| DRD2_HUMAN/371-395 | -ATQMLAIVLGVFIIC**W**LPFFITHIL |
| HRH1_HUMAN/413-437 | -AAKQLGFIMAAFILC**W**IPYFIFFMV |
| OPRD_HUMAN/259-283 | -ITRMVLVVVGAFVVC**W**APIHIFVIV |
| OPRM_MOUSE/278-302 | -ITRMVLVVVAVFIVC**W**TPIHIYVII |
| OPRX_HUMAN/261-285 | -ITRLVLVVVAVFVGC**W**TPVQVFVLA |
| CCR2_HUMAN/241-265 | -AVRVIFTIMIVYFLF**W**TPYNIVILL |
| CCR5_HUMAN/233-257 | -AVRLIFTIMIVYFLF**W**APYNIVLLL |
| AA1R_HUMAN/232-256 | -IAKSLALILFLFALS**W**LPLHILNCI |
| AA2AR_HUMAN/231-255 | -AAKSLAIIVGLFALC**W**LPLHIINCF |
| OPSD_HUMAN/250-274 | -VTRMVIIMVIAFLIC**W**VPYASVAFY |
| OPSD_TODPA/259-283 | -LAKISIVIVSQFLLS**W**SPYAVVALL |
| PAR1_HUMAN/310-335 | RALFLSAAVFCIFIIC**F**GPTNVLLIA |
| PAR2_HUMAN/284-309 | RAIKLIVTVLAMYLIC**F**TPSNLLLVV |
| ----MRs--- | ----------------------------- |
| Q8RUT8_CHLRE/207-232 | RCRQVVTGMAWLFFVS**W**GMFPILFIL |
| W8VZW3_9FLAO/185-210 | GTDSTITKVFWLMMFA**W**TLYPIAYLV |
| N0DKS8_9FLAO/199-224 | AGQKILSNIWILFLIS**W**TLYPGAYLM |
| BACH_HALS3/191-216 | GTAEIFDTLRVLTVVL**W**LGYPIVWAV |
| Q2S2F8_SALRD/184-209 | RVSTLLGNARLLLLAT**W**GFYPIAYMI |
| S5DM51_9ACTN/153-178 | DQAVAFENIKNLILVG**W**IIYPLGYIA |
| BACS2_NATPH/155-180 | GIKSLYVRLRNLTVIL**W**AIYPFIWLL |
| BACR1_HALC1/178-203 | EVASTFNTLTALVLVL**W**TAYPILWII |
| BACR1_HALMA/170-195 | DTRSTFKTLRNLVTVV**W**LVYPVWWLV |
| BACR1_HALWC/174-199 | DTRSTFNALRNIILVT**W**AIYPVAWLV |
| BACR_HALSA/179-204 | EVASTFKVLRNVTVVL**W**SAYPVVWLI |
| BACR2_HALS2/177-202 | EVRSTFNTLTALVAVL**W**TAYPILWIV |
| BACS1_HALSA/155-180 | EQIGLFNLLKNHIGLL**W**LAYPLVWLF |
| Q93WP2_CHLRE/246-271 | ICRDLVRYLAWLYFCS**W**AMFPVLFLL |
| I4DST7_9EURY/165-190 | EVASLFGRLRNLVIVL**W**LLYPVVWIL |

| **Helix 7/G** | **7.45 7.49 7.53** |
|---|---|
| ---GPCRs--- | |
| ACM1_HUMAN/396-424 | PETLWELGYWLCYV**N**STI**N**PMC**Y**ALCNKA |
| ACM3_RAT/521-549 | PKTYWNLGYWLCYI**N**STV**N**PVC**Y**ALCNKT |
| ADRB1_MELGA/321-348 | PDWLFVFFNWLGYA**N**SAF**N**PII**Y**CR-SPD |
| DRD2_HUMAN/404-432 | PPVLYSAFTWLGYV**N**SAV**N**PII**Y**TTFNIE |
| HRH1_HUMAN/446-474 | NEHLHMFTIWLGYI**N**STL**N**PLI**Y**PLCNEN |
| OPRD_HUMAN/296-324 | VVAALHLCIALGYA**N**SSL**N**PVL**Y**AFLDEN |
| OPRM_MOUSE/314-342 | QTVSWHFCIALGYT**N**SCL**N**PVL**Y**AFLDEN |
| OPRX_HUMAN/297-325 | AVAILRFCTALGYV**N**SCL**N**PIL**Y**AFLDEN |
| CCR2_HUMAN/283-311 | LDQATQVTETLGMT**H**CCI**N**PII**Y**AFVGEK |
| CCR5_HUMAN/275-303 | LDQAMQVTETLGMT**H**CCI**N**PII**Y**AFVGEK |
| AA1R_HUMAN/266-294 | PSILTYIAIFLTHG**N**SAM**N**PIV**Y**AFRIQK |
| AA2AR_HUMAN/266-294 | PLWLMYLAIVLSHT**N**SVV**N**PFI**Y**AYRIRE |
| OPSD_HUMAN/284-312 | GPIFMTIPAFFAKS**A**AIY**N**PVI**Y**IMMNKQ |
| OPSD_TODPA/293-321 | TPYAAQLPVMFAKA**S**AIH**N**PMI**Y**SVSHPK |
| PAR1_HUMAN/349-377 | AYFAYLLCVCVSSI**S**CCI**D**PLI**Y**YYASSE |
| PAR2_HUMAN/322-350 | VYALYIVALCLSTL**N**SCI**D**PFV**Y**YFVSHD |
| ----MRs--- |  |
| Q8RUT8_CHLRE/238-266 | GVLSVYGSTVGHTI**I**DLM**SK**NC**W**GLLGHY |
| W8VZW3_9FLAO/216-244 | NADGVVLRQLLFTI**A**DIS**SK**VI**Y**GLMITY |
| N0DKS8_9FLAO/236-264 | SEDGVMARQLVYTI**A**DVS**SK**VI**Y**GVLLGN |
| BACH_HALS3/223-251 | LVQSVGVTSWAYSV**L**DVF**AK**YV**F**AFILLR |
| Q2S2F8_SALRD/221-249 | TPGTIVALQVGYTI**A**DVL**AK**AG**Y**GVLIYN |
| S5DM51_9ACTN/181-209 | VGDFDAIREVLYTI**A**DII**NK**VG**L**GVLVLQ |
| BACS2_NATPH/186-214 | ALLTPTVDVALIVY**L**DLV**TK**VG**F**GFIALD |
| BACR1_HALC1/209-237 | GVVGLGIETLLFMV**L**DVT**AK**VG**F**GFILLR |
| BACR1_HALMA/201-229 | GLVGIGIETAGFMV**I**DLV**AK**VG**F**GIILLR |
| BACR1_HALWC/205-233 | ALTGLYGETLLFMV**L**DLV**AK**VG**F**GFILLR |
| BACR_HALSA/210-238 | GIVPLNIETLLFMV**L**DVS**AK**VG**F**GLILLR |
| BACR2_HALS2/208-236 | GVVGLGIETLAFMV**L**DVT**AK**VG**F**GFVLLR |
| BACS1_HALSA/186-214 | GEATAAGVALTYVF**L**DVL**AK**VP**Y**VYFFYA |
| Q93WP2_CHLRE/277-305 | GHINQFNSAIAHAI**L**DLA**SK**NA**W**SMMGHF |
| I4DST7_9EURY/197-225 | GILPLYWETAAFMV**L**DLS**AK**VG**F**GVVLLR |
